# PfCAP-H is essential for assembly of condensin I complex and karyokinesis during asexual proliferation of *Plasmodium falciparum*

**DOI:** 10.1101/2024.02.26.582160

**Authors:** Pratima Gurung, James P. McGee, Jeffrey D. Dvorin

**Affiliations:** Division of Infectious Diseases, Boston Children’s Hospital, Boston, M.A; Department of Pediatrics, Harvard Medical School, Boston, M.A

**Keywords:** Plasmodium, mitosis, chromosome, condensin I, karyokinesis

## Abstract

Condensin I is a pentameric complex that regulates the mitotic chromosome assembly in eukaryotes. The kleisin subunit CAP-H of the condensin I complex acts as a linchpin to maintain the structural integrity and loading of this complex on mitotic chromosomes. This complex is present in all eukaryotes and has recently been identified in *Plasmodium spp*. However, how this complex is assembled and whether the kleisin subunit is critical for this complex in these parasites is yet to be explored. To examine the role of PfCAP-H during cell division within erythrocytes, we generated an inducible PfCAP-H knockout parasite. We find that PfCAP-H is dynamically expressed during mitosis with the peak expression at the metaphase plate. PfCAP-H interacts with PfCAP-G and is a non-SMC member of the condensin I complex. Notably, the absence of PfCAP-H does not alter the expression of PfCAP-G but affects its localization at the mitotic chromosomes. While mitotic spindle assembly is intact in PfCAP-H deficient parasites, duplicated centrosomes remain clustered over the mass of unsegmented nuclei with failed karyokinesis. This failure leads to the formation of an abnormal nuclear mass, while cytokinesis occurs normally. Altogether, our data suggest that PfCAP-H plays a crucial role in maintaining the structural integrity of the condensin I complex on the mitotic chromosomes and is essential for the asexual development of malarial parasites.

**Importance:** Mitosis is a fundamental process for *Plasmodium* parasites, which plays a vital role in their survival within two distinct hosts - human and *Anopheles* mosquitoes. Despite its great significance, our comprehension of mitosis and its regulation remains limited. In eukaryotes, mitosis is regulated by one of the pivotal complexes known as condensin complexes. The condensin complexes are responsible for chromosome condensation, ensuring the faithful distribution of genetic material to daughter cells. While condensin complexes have recently been identified in *Plasmodium spp*, our understanding of how this complex is assembled and their precise functions during the blood stage development of *Plasmodium falciparum* remains largely unexplored. In this study, we investigate the role of a central protein, PfCAP-H, during the blood stage development of *P. falciparum*. Our findings reveal that PfCAP-H is essential and plays a pivotal role in upholding the structure of condensin I and facilitating karyokinesis.

## Introduction

*Plasmodium falciparum* is a protozoan parasite that is responsible for the most severe forms of human malaria. Malaria remains one of the most important global infectious diseases, claiming more than 619 000 lives worldwide annually [1]. These parasites have a complex life cycle that alternates between two different hosts, *Anopheles* mosquitoes and humans. To survive and thrive in these hosts, they replicate through an atypical cell division. These parasites follow schizogony within hepatocytes and red blood cells (erythrocytes) in their human hosts. On the other hand, they undergo gametogenesis and sporogony in mosquitoes [2]. Among all these stages, the extensive proliferation of parasites within human erythrocytes causes the signs and symptoms of clinical malaria. Parasite cell division in erythrocytes, known as schizogony, is an unconventional mode of cell division that includes growth and budding phases [3, 4]. During the growth phase, parasite undergo several asynchronous rounds of DNA replication and mitosis (S–M phase) without cytokinesis [5, 6]. Asexual *Plasmodium* parasites undergo closed mitosis, where decondensed chromosomes are segregated, followed by nuclear division with an intact nuclear envelope throughout the cycle [2]. Later in the budding phase, the parasite undergoes a final round semi-synchronous nuclear division along with cytokinesis to produce mature daughter cells, called merozoites. These merozoites invade new erythrocytes to begin the proliferation cycle again [7, 8]. Despite its predominant role in asexual proliferation, the regulation of this atypical mitosis is still underexplored in *Plasmodium* parasites.

During mitosis, chromatin condensation and segregation are important events to ensure that genomic material is equally divided into the daughter nuclei [9, 10]. This process is mediated by two distinct condensin complexes, condensin I and II. These complexes are pentameric and comprised of two parts – core subunits and regulatory subunits. The core subunit consists of structural maintenance of chromosomes (SMC) 2 and 4, common in condensin I and II complexes. In contrast, the regulatory subunits, collectively known as chromosome-associated proteins (CAP) or non-SMC members, differ between two complexes; condensin I contains CAP-H/CAP-G/CAP-D2 while condensin II has CAP-H2/CAP-G2/CAP-D3 [11, 12]. While condensin I is highly conserved across eukaryotes and its localization is dynamic throughout the cell division, condensin II remains in nucleus and is absent in some organisms (e.g., yeast and insects) [12, 13].

In *Plasmodium spp.*, both condensin I and II complex have been identified by homology prediction [14, 15]. The two core subunits (SMC 2/4) and one non-SMC member-PfCAP-G of the condensin I complex have been genetically interrogated in *Plasmodium spp.* [14, 16]. In *Plasmodium berghei,* SMC2/4 displayed a dynamic localization in both asexual parasites and gametocytes. PbSMC2/4 knockout in asexual parasites was unsuccessful, therefore a functional evaluation was not possible during that stage. Depletion of PbSMC2/4 during gametocytogenesis impaired male gametogenesis and zygote differentiation and thus blocked parasites transmission in *P. berghei* [14].

On the other hand, the knockdown of PfCAP-G (or Merozoite Organizing Protein - MOP), a non-SMC member of the condensin I complex, showed a fitness defect in the asexual development of *P. falciparum*. The PfCAP-G-deficient parasites showed flawed segmentation with a large residual agglomerate of partially divided cells [16]. Given that the PfCAP-G knockdown phenotype was incomplete, likely due to insufficient protein knockdown, the phenotype of complete loss of condensin I remains insufficiently evaluated. Furthermore, there remains a lack of experimental evidence on how these complexes are assembled on chromosomes and their function as regulators of mitosis during the asexual blood stage development in *P. falciparum*.

In eukaryotes, the kleisin subunit CAP-H acts as a linchpin in the assembly of the condensin I complex [17]. The CAP-H sequence consists of five motifs that bind to different components of the condensin I complex [18–20]. The N- and C-terminal motifs of this protein interact with the core proteins (SMC 2 and SMC 4), while the central regions consist of motifs that bind to two other non-SMC proteins-CAP-G and CAP-D2 along with a region that interacts with chromosomal DNA to anchor condensin [18, 21–23]. In addition, the loading of the condensin I complex during mitosis is regulated by the N-terminal tail of CAP-H in *Xenopus* egg extracts [24]. Mutation or deletion of CAP-H results in a mitotic chromosome condensation and segregation defect in yeast and *Drosophila melanogaster* [25–27].

CAP-H has been bioinformatically predicted in *Plasmodium spp.* [14], and we have focused on interrogating its function during the asexual development of *Plasmodium falciparum* within human erythrocytes. We generated a parasite strain allowing inducible knockout of PfCAP-H. We show that PfCAP-H is a member of the condensin I complex and is essential for the asexual parasite replication. PfCAP-H is dynamically localized during mitosis and can be used as a marker for the metaphase plate. Depletion of PfCAP-H causes abnormal karyokinesis, while cytokinesis occurs normally. This study provides new insights into the function of the condensin I complex during asexual replication of *P. falciparum*.

## Results

### PfCAP-H has conserved N- and C-terminal region and is expressed in proliferative blood stages

CAP-H is a central component of the condensin I complex in all eukaryotes [17]. Bioinformatic analysis predicted that CAP-H is also present in *Plasmodium falciparum* [14]. PF3D7_1304000 (hereafter referred to as PfCAP-H) is 1024 amino acids long with a putative condensin complex subunit 2 domain. To comprehensively evaluate the PfCAP-H sequence by *in silico* analysis, we compared the sequence of PfCAP-H with CAP-H homologs from a wide range of eukaryotes. PfCAP-H consists of conserved regions in its N (1 – 250 aa) and C (890 – 1000 aa) termini, with a less conserved interior sequence (Fig. S1A). The complete amino acid sequence has 21-34% similarity with the majority of the non-Apicomplexan homologs and, as expected, higher similarity to *Plasmodium* homologs (e.g., 64% to PbCAP-H) (Fig. S1B).

To directly investigate the role of PfCAP-H in *Plasmodium falciparum*, we generated an inducible PfCAP-H knockout (iKO) strain in 3D7*pfs47*DiCre parasites (named PfCAP-H^DiCre^) [28] (Fig. 1A). In these parasites, the native PfCAP-H gene locus has been replaced with a *loxP*-flanked, codon-altered PfCAP-H with the spaghetti monster (sm)V5 epitope tag at the C-terminus. Integration of the donor cassette was verified by PCR amplification. The expected size of 2.3 and 0.98 kilobase-pairs (kb) was observed in the transgenic line, while no equivalent bands were amplified in the parental line (Fig. S2A).

**Figure 1:**
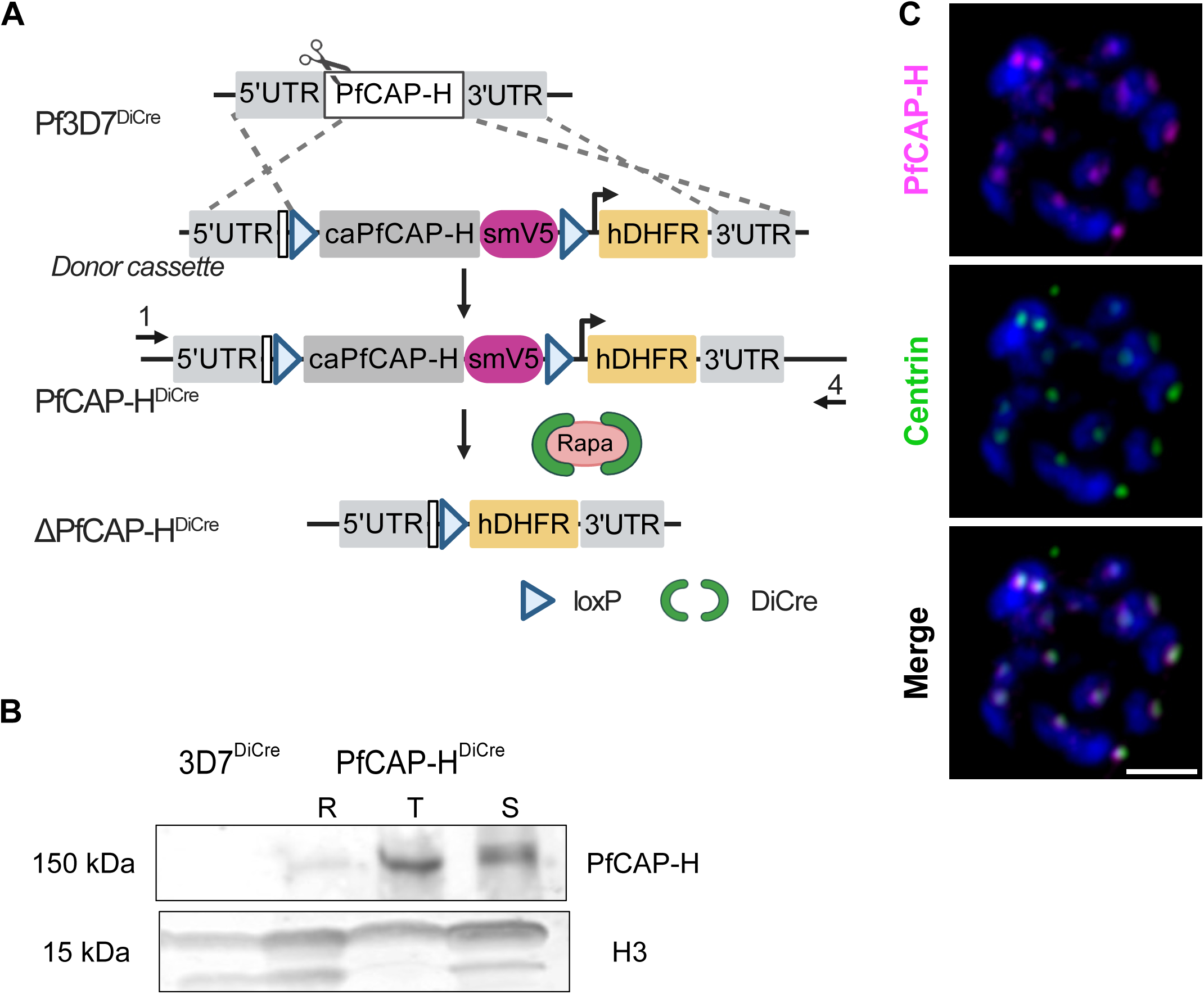
Expression and localization of PfCAP-H in PfCAP-H^DiCre^ parasites. **(A)** Schematic of PfCAP-H^DiCre^ parasites. In PfCAP-H^DiCre^ parasites, we replaced the endogenous locus of *PfCAP-H* in 3D7*pfs47*DiCre (referred to as 3D7^DiCre^) parasites with a loxP-flanked (triangle), codon-altered PfCAP-H with spaghetti monster (sm)-V5 tag (magenta-colored box) at the C-terminus. **(B)** Immunoblot showing the expression of PfCAP-H during asexual blood stages (R - Rings, T - Trophozoites, S - Schizonts) probed with α-V5. The parental Pf3D7^DiCre^ parasites are used as a negative control and α-Histone H3 was used as a protein loading control. **(C)** The localization of PfCAP-H was visualized by α-V5 (magenta) for smV5 tagged PfCAP-H and α*-*Centrin (green) was used as a marker of centrosome by slide-based IFA. The IFA showed that PfCAP-H is localized near centrosome. The DNA was stained with Hoechst 33342 (blue). Scale bar = 2 µm.

We next sought to determine the expression timing and localization of PfCAP-H during the asexual blood stages. The immunoblot, probed with α-V5, demonstrated the expected 150 kDa of smV5-tagged PfCAP-H in both trophozoites and schizonts (Fig. 1B). The observed PfCAP-H expression agrees with the transcriptional profiling data [15], indicating a likely role for PfCAP-H during asexual development. Given that PfCAP-H homologs are present near centrosomes in other organisms [12], we performed an immunofluorescence assay (IFA) on schizonts with an antibody that recognizes *Plasmodium* centrins as a marker for the centrosome in these parasites. PfCAP-H localizes near the centrosomes (Fig. 1C), similar to the reported localizations for PbSMC2/4 [14].

### PfCAP-H is highly expressed in mitotically active nuclei during schizogony

To interrogate the subcellular localization of PfCAP-H during the proliferative stages of asexual development, we performed IFAs throughout the schizont stage of the asexual development cycle, probing for PfCAP-H (α-V5) together with the nuclear (DNA) stain Hoechst 33342. The IFA revealed discrete perinuclear foci throughout schizogony (Fig. S3), with diminished protein detection when segmentation is complete. This expression resembles the pattern of proteins crucial for cell cycle progression [29].

For higher resolution imaging, we used ultrastructure-expansion microscopy (U-ExM) to more precisely localize PfCAP-H and examine its role during cell division in these parasites. For these studies, we have included a fluorophore conjugated to N-hydroxysuccinimide (NHS) (herein referred to as “NHS ester”) as a non-specific stain for protein density [30], SYTOX Deep Red as a DNA stain, α-V5 for smV5-tagged PfCAP-H, and α-tubulin for mitotic spindles. As noted above, *P. falciparum* undergoes an atypical cell division known as schizogony, where individual nuclei undergo asynchronous S/M cycles, followed by a final semi-synchronous cycle coupled with the budding of the daughter cells [4, 29]. Given the complexity of schizogony and decondensed chromosomes in *Plasmodium falciparum*, it is challenging to distinctly visualize different stages of mitosis in these parasites. Therefore, we relied upon the positioning of mitotic apparatus/microtubule organizing centers and microtubules to evaluate the distinct phases of mitosis in these parasites. As shown in the Fig. 2A, in nuclei undergoing mitosis, we observed that PfCAP-H shows a speckled pattern at the plus end tip of the mitotic spindles. The signal intensifies and organizes as clusters at the metaphase plate where chromosomes are typically aligned during metaphase. Subsequently, these signals resume their speckled pattern at the tip of mitotic spindles during anaphase and telophase, which finally becomes more diffuse at the end of the cell cycle. To further confirm the localization of PfCAP-H to the plus end of mitotic spindles, we generated a parasite strain where the kinetochore marker, PfNDC80/PF3D7_0616200[31, 32], was additionally epitope-tagged at its endogenous locus with spaghetti monster HA (smHA) (Fig. S4A and S4B). Immunofluorescence of these dual-tagged parasites demonstrated that PfCAP-H localizes within the surrounding PfNDC80 staining, suggesting that PfCAP-H is on the chromosomal side (Fig. S4C). These results demonstrate that PfCAP-H shows a dynamic pattern during different stages of mitosis and provides a marker for the metaphase plate in mitotically active parasites.

**Figure 2:**
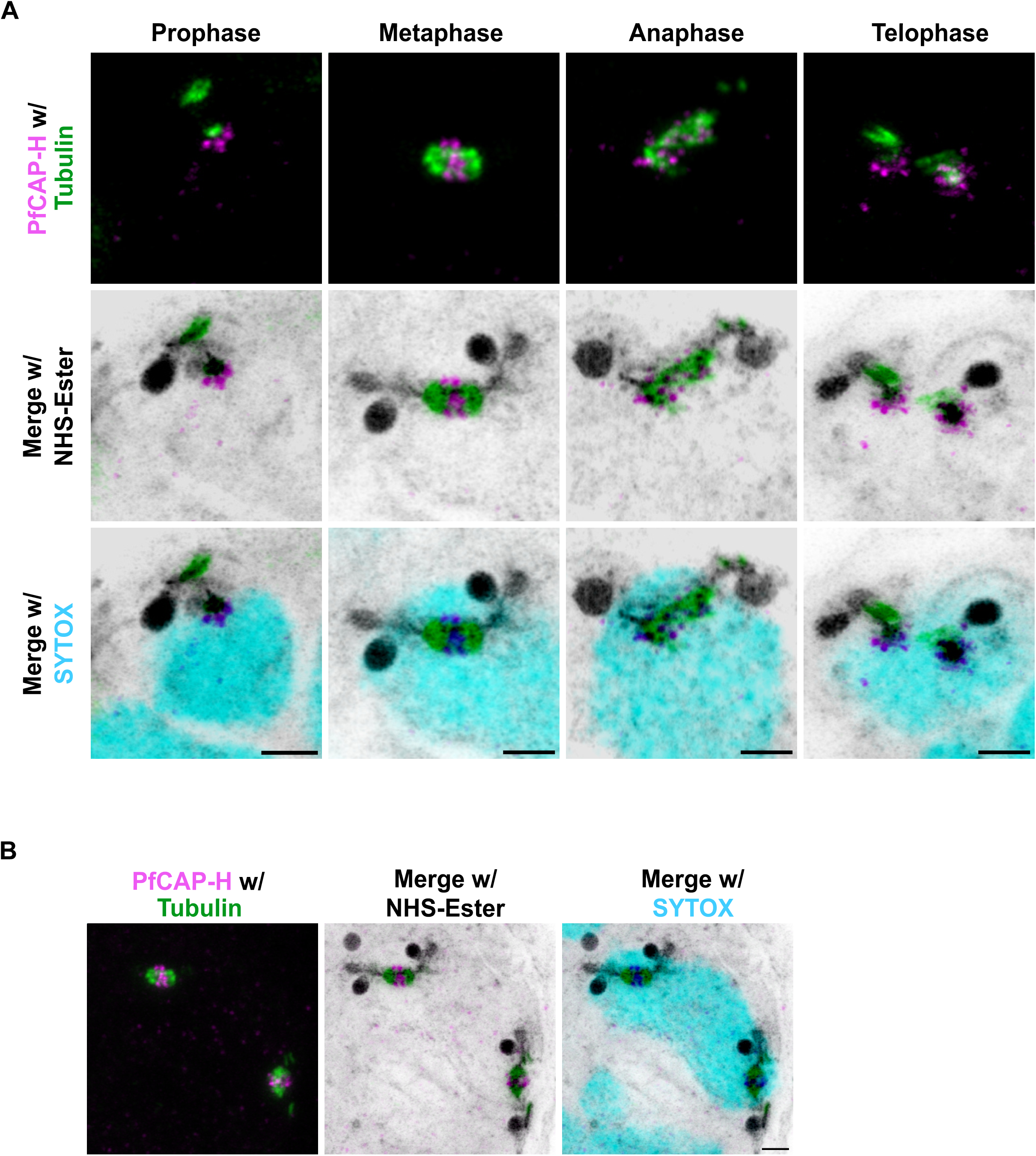
PfCAP-H is present in mitotically active nuclei. **(A)** PfCAP-H^DiCre^ parasites were prepared for U-ExM and stained with α-V5 for PfCAP-H (in magenta), α-tubulin for mitotic spindles (in green), NHS-Ester for protein stain (in grayscale), and SYTOX for nuclear stain (in cyan) and captured using Airyscan microscopy. With the cues from the localization of mitotic spindles, different stages of mitosis (prophase, metaphase, anaphase, and telophase) are marked compared to position of mitotic spindles in conventional mitotic stages. The peak expression of PfCAP-H is at the metaphase plate. Scale bars = 2 µm. **(B)** Example of schizont exhibiting two mitotic spindles within one dividing nuclei and thus suggesting the existence of schizogony with limited karyokinesis during blood stages. Scale bars = 2 µm, image is projection of 20 z-slices.

Interestingly, we observed that, in ∼10% of schizonts (3 of 31), two duplicated hemi-spindle and/or mitotic spindles are present within a single nucleus (Fig. 2B). These unusual events suggest that these schizonts follow unconventional schizogony where already duplicated genomes with a single, undivided nucleus have already started the next round of mitosis before completing karyokinesis. This supports a model of schizogony with limited karyokinesis [4] occurring at times during asexual development.

### PfCAP-H is a member of the condensin I complex in *Plasmodium falciparum*

The observed expression and localization pattern of PfCAP-H resembles the pattern exhibited by SMC core members (PbSMC2/4) and PfCAP-G of the condensin I complex in *Plasmodium spp*. [14, 16], suggesting that PfCAP-H is a member of the condensin I complex in these parasites. To confirm this hypothesis in *Plasmodium falciparum*, we performed two independent experiments. First, we investigated whether PfCAP-H interacts with PfCAP-G, a non-SMC member of the condensin I complex by performing IFA with PfCAP-H^DiCre^ /PfCAP-G parasites, where the endogenous PfCAP-G has a smHA epitope in the smV5-tagged PfCAP-H strain. The dual-transgenic line was confirmed by integration PCR and whole genome sequencing (Fig. S5A-S5C, sequence reads deposited in NCBI Sequence Read Archive, #XXXXXX). PfCAP-H and PfCAP-G colocalize with each other and displayed a similar dynamic expression pattern throughout schizogony (Fig. 3A and S5D), suggesting that these two proteins interact with each other throughout this stage. Second, we used an alternative approach where we fused a promiscuous version of the biotin ligase BirA[33] to PfCAP-G in 3D7 parasites to generate a new transgenic PfCAP-G^BirA^ strain. We used this PfCAP-G^BirA^ parasites to identify the proteins that interact with PfCAP-G using BioID-based proximity labeling in late schizonts coupled with mass spectrometry (Fig. 3B and Supplemental Table S1) [33]. As anticipated, all the members of the condensin I complex including PfCAP-H were among the top hits, implying that PfCAP-G is a true partner of PfCAP-H along with other members of the condensin I complex. Altogether, these data establish PfCAP-H as a *bona fide* member of the condensin I complex in *P. falciparum*.

**Figure 3:**
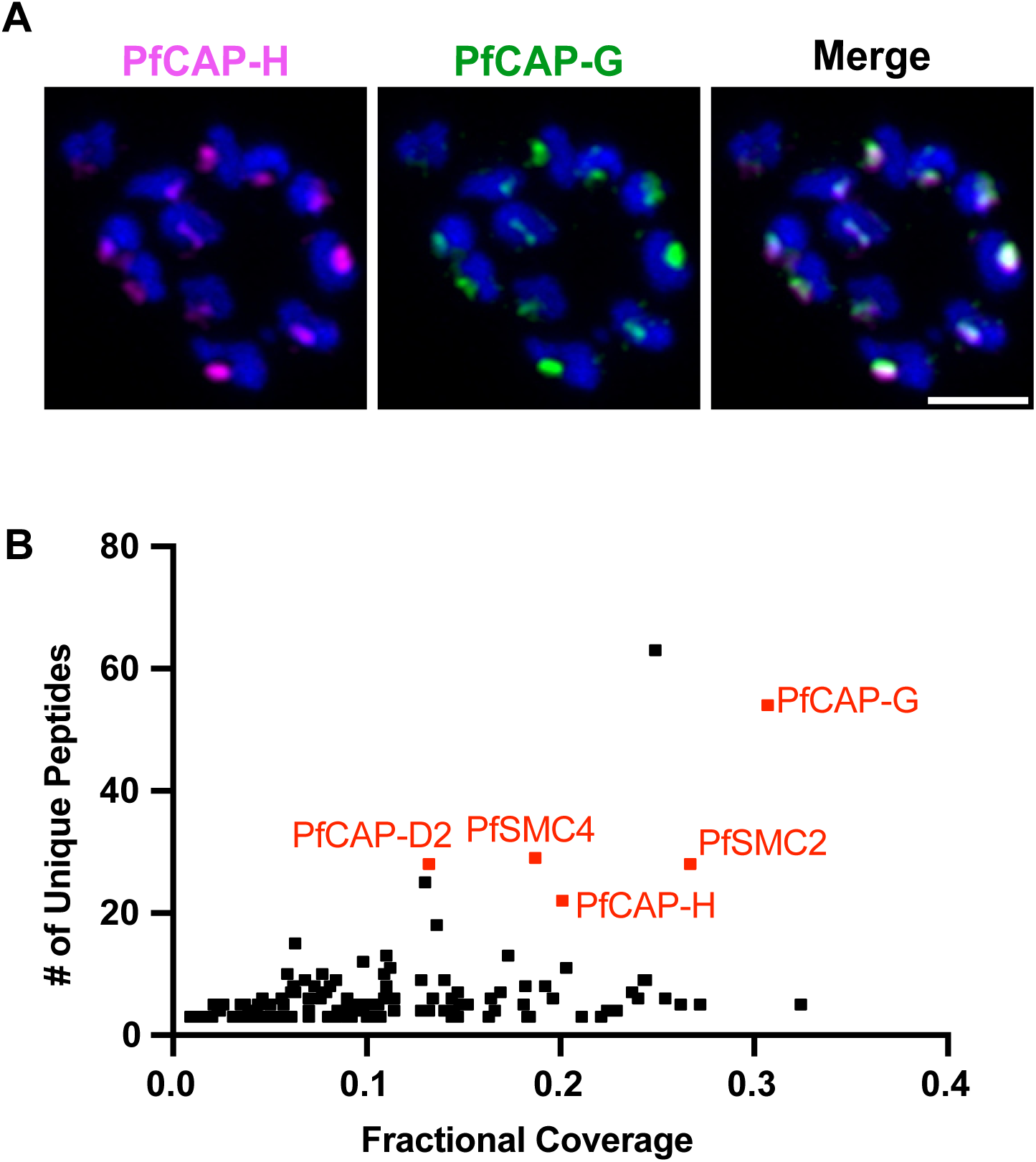
PfCAP-H is a member of the condensin I complex. **(A)** Schizonts from PfCAP-H^DiCre^/PfCAP-G parasites were fixed and stained with α-V5 against PfCAP-H and α-HA against PfCAP-G (n= 3). The slide-based IFA showed that PfCAP-H colocalizes with PfCAP-G. Scale bars = 2 µm **(B)** Proximity labeling in PfCAP-G^BirA^ *parasites* demonstrates that PfCAP-H interacts with PfCAP-G and other members of the condensin I complex (representative example shown from one of three biological replicates).

### PfCAP-H is essential for asexual development of blood stage parasites

The condensin I complex plays a crucial role during cell division. Knockdown of PfCAP-G produces a significant fitness defect in *P. falciparum*, and PbSMC2/4 could not be knocked out in *P. berghei* asexual parasites[14, 16]. To directly determine the consequences for complete removal of PfCAP-H, we performed a replication growth assay where we treated synchronized early ring (0-4 hrs) staged PfCAP-H^DiCre^ parasites with rapamycin or DMSO and monitored their growth for two consecutive cycles using flow cytometry. In these parasites (Fig. 1A), rapamycin dimerization of the DiCre subunits mediates excision of the PfCAP-H gene flanked by two *loxP* sites, resulting in a PfCAP-H knockout (PfCAP-H iKO) [28]. In PfCAP-H iKO parasites, whole locus PCR amplification detected 4.8 kb truncated gene locus in rapamycin treated parasites compared to full length 9.5 kb modified PfCAP-H locus in DMSO-treated parasites (Fig. S2A). Furthermore, IFA demonstrated that the rapamycin treatment resulted in complete loss of PfCAP-H protein in PfCAP-H iKO parasites (Fig. S2B) We used the parental 3D7*pfs47*^DiCre^ parasite strain as a control. The rapamycin-treated PfCAP-H^DiCre^ parasites exhibited a 97.8% ± 0.6 growth defect resulting in death of parasites within the same cycle of treatment when compared to DMSO-treated parasites (Fig. 4A).

**Figure 4:**
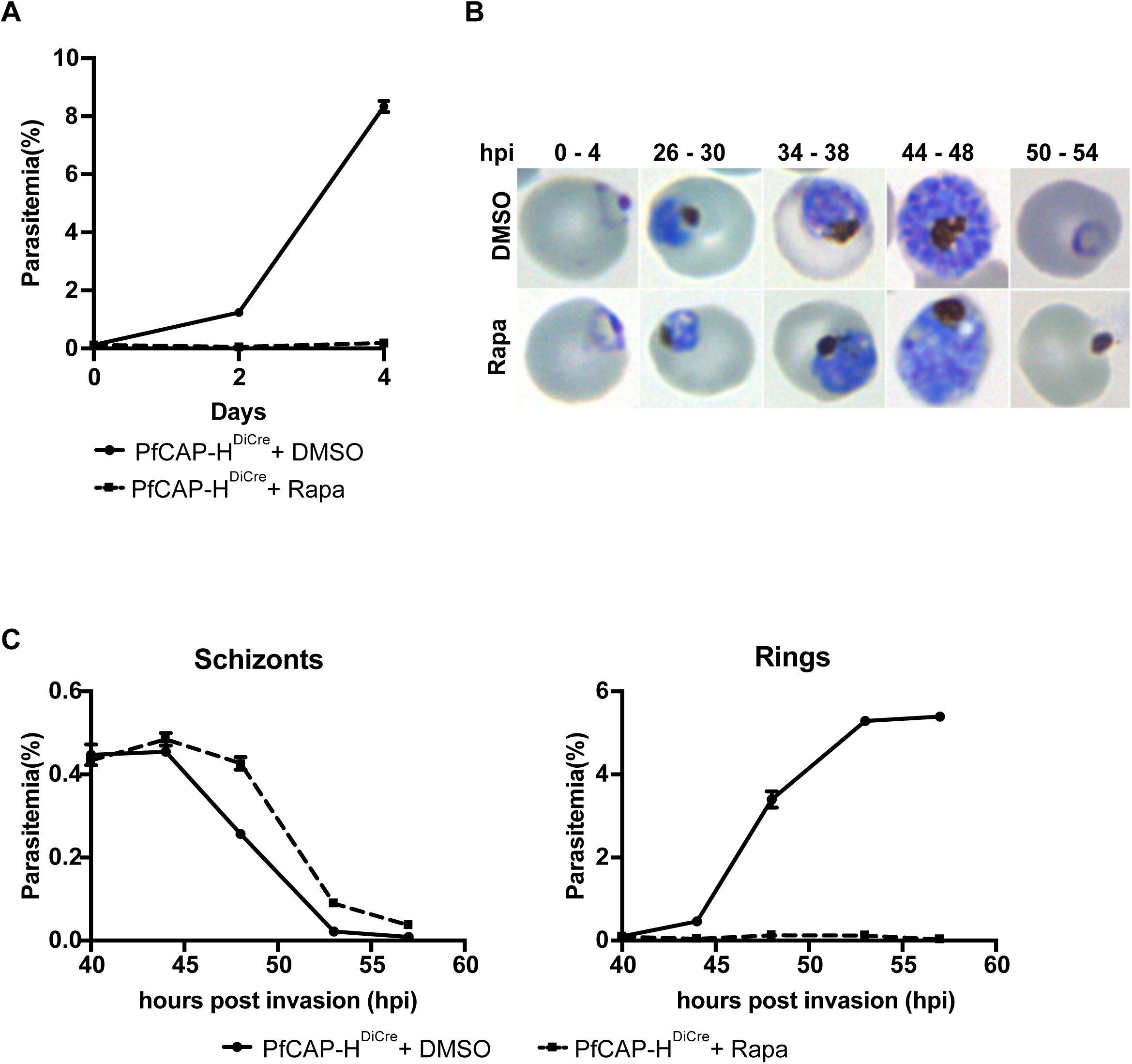
PfCAP-H is essential for asexual development of parasites. **(A)** PfCAP-H^DiCre^ parasites were treated with rapamycin and their growth was monitored over two consecutive cycles by flow cytometry. Loss of PfCAP-H showed a lethal effect on parasite asexual growth development. Error bars show standard deviation from mean of three independent biological replicates. **(B)** The Hemacolor stained smear was prepared for the first growth cycle to observe progression of asexual stages. In rapamycin treated parasites, we observed that the parasites are arrested at the late blood stages and no new invasion of red blood cells were observed. **(C)** Time course of PfCAP-H parasites cultured with rapamycin/DMSO were collected every 5 hours from 40 to 60 hpi, and parasitemia was measured by flow cytometry (n = 3, error = standard error of mean (SEM)) to examine the duration of egress and invasion. Rapamycin-treated parasites did not show any significant difference in the progression of growth, however they could not invade new red blood cells.

To evaluate the growth phenotype in more detail, we examined field-stained parasites to determine the developmental stage of the arrested parasites. At this level of resolution, we observed that PfCAP-H iKO parasites grew normally until the early schizont stage (approximately 30 hours post-invasion [hpi]). After this point, parasites had abnormal morphology and, eventually, failed to generate newly invaded rings (Fig. 4B). Because we did not observe an accumulation of unruptured schizonts, we measured the timing of egress (and potential reinvasion or lack thereof), in the presence and absence of PfCAP-H knockout, by flow cytometry. Notably, we did not find any significant difference in the timing of schizont egress in the absence of PfCAP-H. PfCAP-H iKO parasites ruptured at the same time as control parasites, suggesting that loss of PfCAP-H does not cause an egress defect. However, the egressed PfCAP-H iKO parasites did not form new rings (Fig. 4C).

We next asked whether this growth defect is due to alterations in DNA replication in PfCAP-H iKO parasites. To address this, we measured the DNA content of rapamycin or DMSO treated parasites at 40 hpi, when parasites are at the peak of their dividing state. Parasite DNA was labeled with SYBR Green and quantified by measurement of mean fluorescence intensity (MFI) by flow cytometry. Remarkably, we did not find any significant difference in the total DNA content in PfCAP-H iKO and control parasites (Fig. S6), indicating that deletion of PfCAP-H does not have a major effect on DNA replication in the knockout parasites.

### PfCAP-H depletion does not affect the centrosome and mitotic spindle formation

Centrosomes act as the microtubule-organizing center (MTOC), and failure to duplicate or separate could lead to aberrant cell division [34]. We interrogated whether the depletion of PfCAP-H affects the biogenesis and dynamics of centrosomes during mitosis with U-ExM on the DMSO and rapamycin-treated PfCAP-H^DiCre^ strains probed with α-V5, α-Centrin, NHS-Ester, and SYTOX [35]. Surprisingly, the centrosomes are present and duplicated normally, even without PfCAP-H (Fig. 5A and 5B). However, in the absence of PfCAP-H, the centrosomes remain clustered over the mass of unsegmented nuclei. These results suggest that the phenotype observed by depletion of PfCAP-H is not due to failure in centrosome duplication or separation but to some other defect.

**Figure 5:**
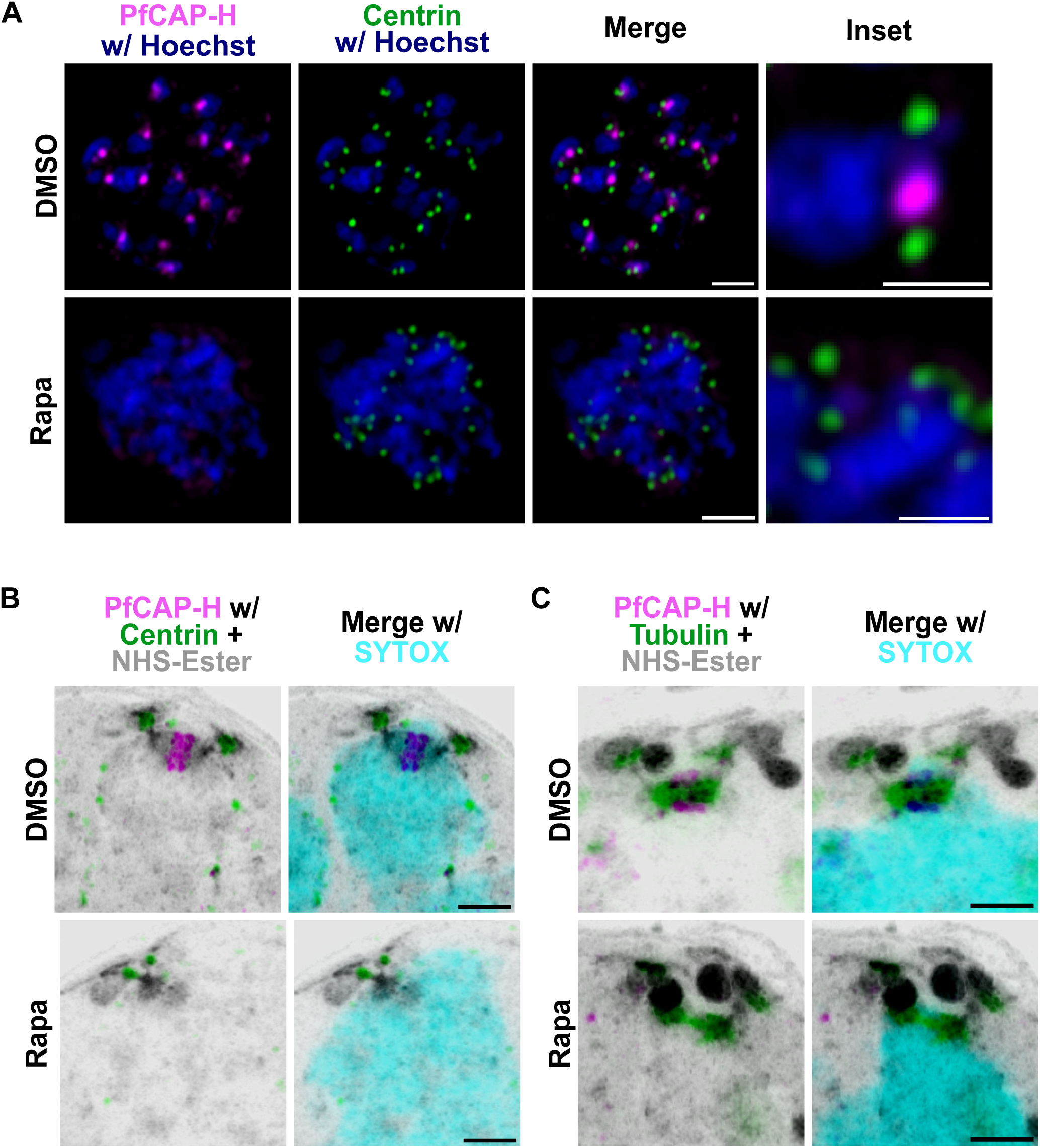
PfCAP-H is dispensable for centrosome dynamics and microtubule formation. PfCAP-H^DiCre^/PfCAP-G parasites were synchronized, treated with rapamycin/DMSO, and collected for slide-based IFA and U-ExM. The samples were stained with α-V5 (PfCAP-H, magenta), α-Centrin (Centrosome marker, green), and α-tubulin (green). In addition, for U-ExM, NHS-Ester (greyscale) and SYTOX (cyan) was used. **(A)** In PfCAP-H deficient parasites, the slide-based IFA demonstrated that centrosome is duplicated but their separation is affected. The highly resolved U-ExM further confirms the similar observation. The U-ExM showed that centrosome **(B)** and microtubule formation **(C)** look similar in both the conditions. Scale bar = 2 µm for images and inset.

We monitored if mitotic spindles could form in the absence of PfCAP-H by direct visualization of these structures by U-ExM with α-tubulin in dividing parasites. In rapamycin-treated parasites, we observed normal mitotic spindle formation, suggesting that the PfCAP-H is also not required for mitotic spindle assembly (Fig. 5C). Taken together, these results implies that the PfCAP-H function is dispensable for centrosome and mitotic spindle assembly.

### Loss of PfCAP-H deters the proper localization of PfCAP-G on mitotic chromatin

PfCAP-H homologs play a vital role in maintaining the ring-like structure of condensin I complex and ensuring their collective function as chromatin condensers in other eukaryotes [17]. To investigate if this function was preserved for PfCAP-H, we evaluated the expression and localization of PfCAP-G in the presence or absence of PfCAP-H. With U-ExM, we found that the PfCAP-G is no longer localized to the mitotic chromosomes in the PfCAP-H iKO parasites (Fig. 6A). However, the expression of PfCAP-G is not affected in the absence of PfCAP-H (Fig. 6B). This observation suggests that the depletion of PfCAP-H does not affect the expression of PfCAP-G, but it is crucial for the loading of PfCAP-G on the mitotic chromosomes. Thus, the data implies that PfCAP-H is critical for the assembly for non-SMC members of the condensin I complex in *Plasmodium falciparum*.

**Figure 6:**
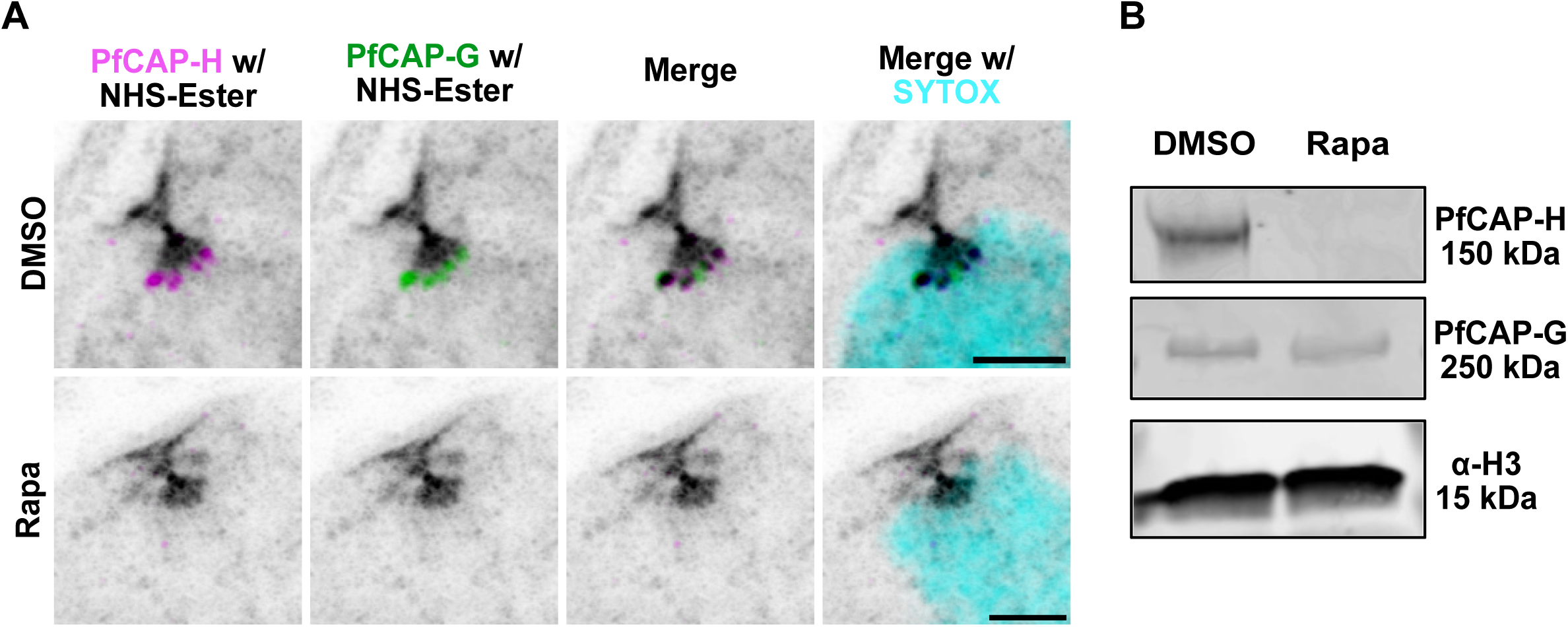
PfCAP-H is essential for assembly of condensin I complex. PfCAP-H^DiCre^/PfCAP-G parasites were synchronized, treated with rapamycin/DMSO, and collected for U-ExM. The samples were stained with α-V5 (PfCAP-H, magenta), and α-HA (PfCAP-G, green), NHS-Ester (greyscale), and SYTOX (cyan). **(A)** In PfCAP-H deficient parasites, the highly resolved U-ExM showed that localization of PfCAP-G to the mitotic chromosome is lost in the absence of PfCAP-H. **(B)** The immunoblot analysis displayed that in PfCAP-H KO parasites expression of PfCAP-G is unaffected. Scale bar = 2 µm.

### Depletion of PfCAP-H affects karyokinesis but does not impede cytokinesis

Our observations thus far suggest that the defect in PfCAP-H iKO parasites is due to abnormal nuclear division rather than DNA replication. We thus interrogated the process of karyokinesis in these parasites during their asexual development. Again, we utilized U-ExM along with NHS-Ester, α-V5 for PfCAP-H, and SYTOX to visualize the parasite DNA. At 40 hpi we observed that PfCAP-H iKO parasites showed defective nuclear division. At the same stage, the control parasites had normal nuclear division (Fig. S7). We further followed the karyokinesis until the end of one development cycle by trapping parasites prior to egress with the cysteine protease inhibitor E64 [36]. Strikingly, we observed that fully matured PfCAP-H iKO parasites showed a large agglomerate of incompletely separated nuclear material compared to the >20 normally separated nuclei in control parasites (Fig. 7A). This result suggests that the defect of chromosome segregation leads to a defect in karyokinesis.

**Figure 7:**
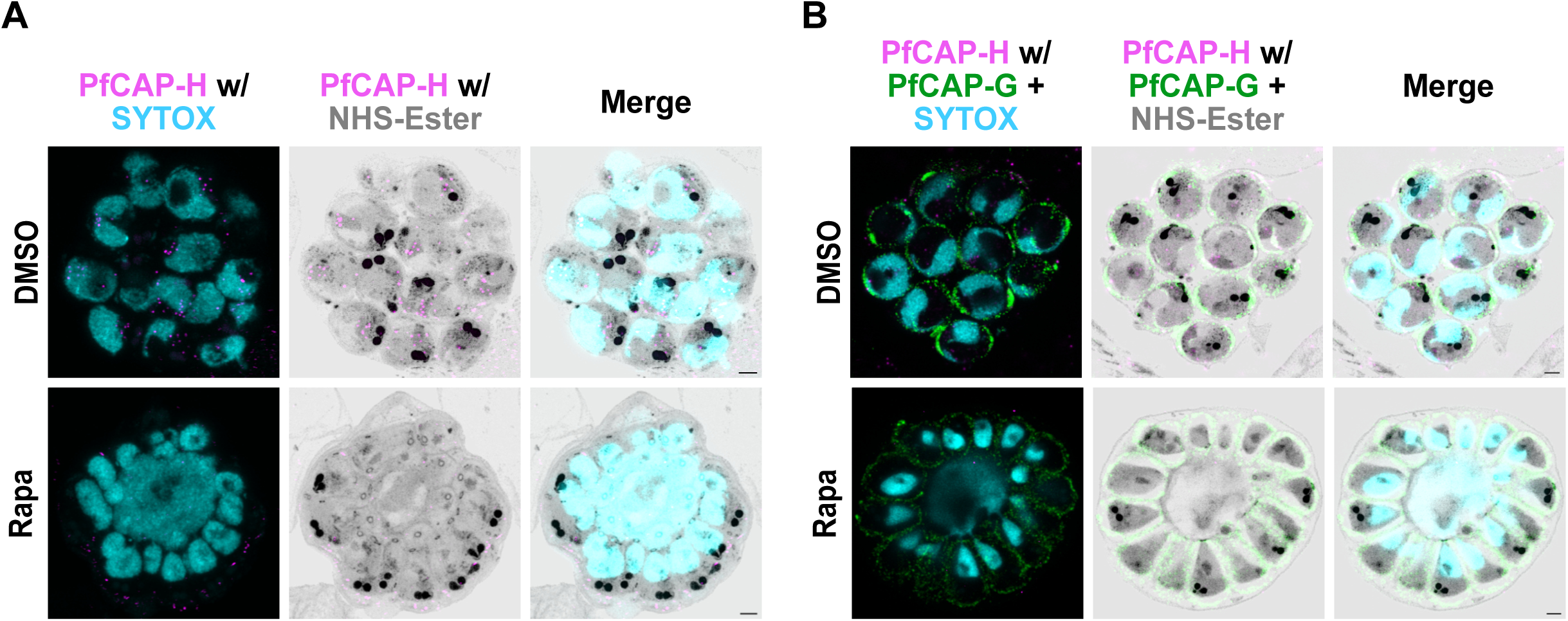
PfCAP-H knockout affects karyokinesis but not cytokinesis. Synchronized PfCAP-H^DiCre^ parasites were treated with rapamycin or DMSO, probed with α-V5 (PfCAP-H, magenta), α-PfGAP45 (IMC, green), NHS-Ester (protein, greyscale), and SYTOX (nucleus, cyan) for U-ExM (n= 3). **(A and B)** E64-arrested late matured schizonts, PfCAP-H KO parasites showed giant agglomerate of incompletely separated nuclei while the IMC appeared to be surrounding all potential merozoites, implying unaffected cytokinesis. Scale bar = 2 µm.

To evaluate cytokinesis, E64-treated parasites were stained with antibodies against V5 and PfGAP45, an inner membrane complex (IMC) associated protein.[37] With U-ExM, we demonstrated that despite abnormal karyokinesis, the IMC still surrounds the nascent merozoites. A varied amount of nuclear material is contained in the forming merozoites with some nuclear material observed in streaks within the contracted basal complex as well as additional nuclear material in the residual body (Fig. 7B). While there may be abnormalities in the final steps of abscission, these results suggest that the processes of cytokinesis, including IMC formation and basal complex contraction, are largely intact. These schizonts still egress (Fig. 4C) but do not form new ring-stage parasites.

### PfCAP-H is likely dispensable for sexual stage development

Given that PbSMC2/4 and PfCAP-G are necessary for the sexual development of these parasites, we asked whether PfCAP-H is required for the sexual development of *Plasmodium falciparum*. To investigate the role of PfCAP-H in gametocyte development, we induced sexual commitment in synchronized PfCAP-H^DiCre^ parasites, treated with rapamycin or DMSO, and then monitored the gametocytogenesis over 12 days. The gametocyte conversion rate was similar in both PfCAP-H iKO and control parasites. In addition, the absolute gametocytemia on day 6 after induction was similar at 1.9 +/- 0.3% and 2.0 +/- 0.1% (mean +/- SD) in the DMSO and rapamycin conditions, respectively. In stage III gametocytes, PfCAP-H localizes adjacent to the nucleus (Fig. 8A). Intriguingly, we did not find any significant morphological difference between PfCAP-H iKO and control parasites throughout the 12 days of the assay (Fig. 8B, S8A, and S8B).

**Figure 8:**
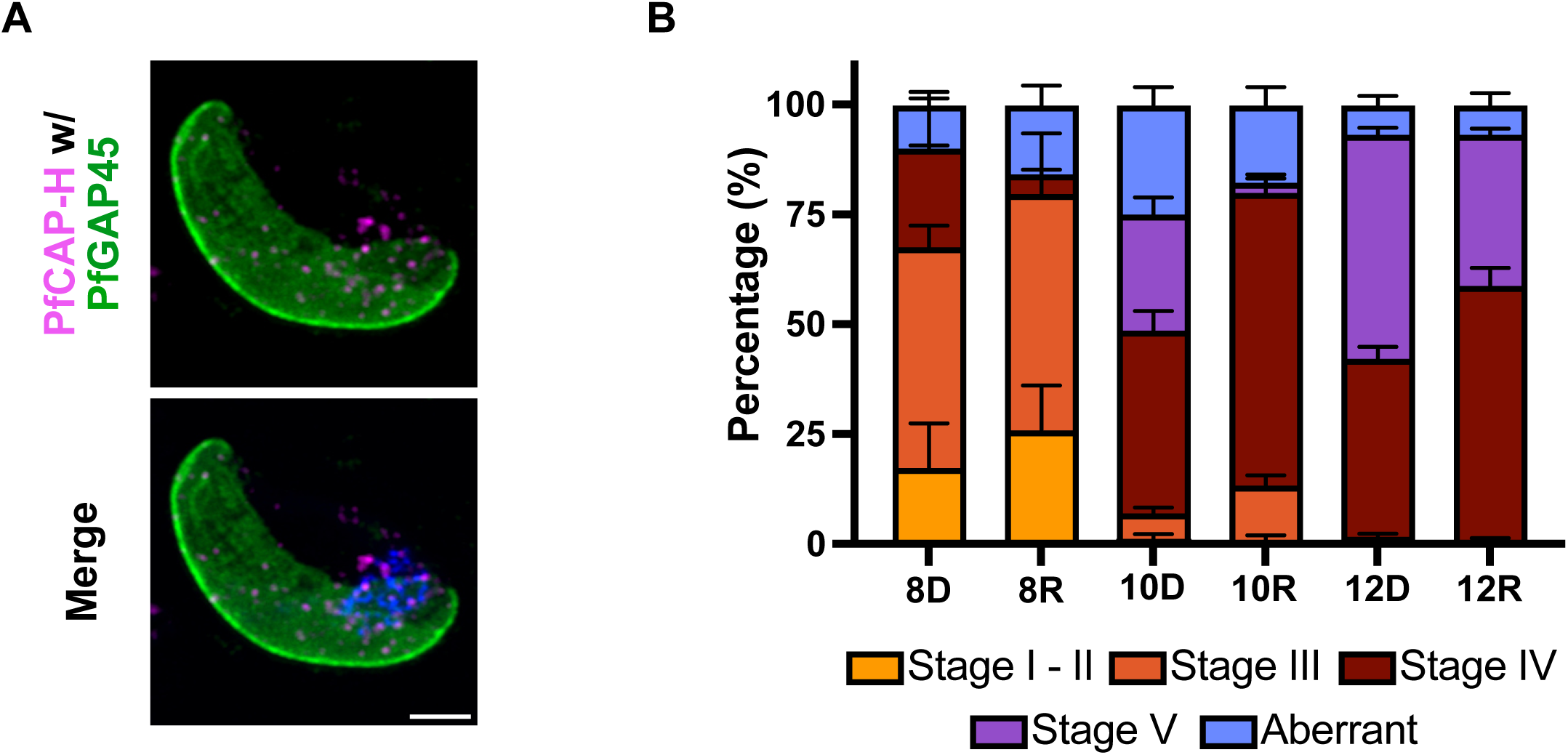
PfCAP-H is likely dispensable for gametogenesis. **(A)** Slide-based IFA was performed for the localization of PfCAP-H in sexual stages (Stage III) on Day 8 post-induction. The slides were probed with α-V5 (magenta) for PfCAP-H and α-PfGAP-45 as IMC markers (green). IFA showed that PfCAP-H is present adjacent to the nucleus in stage III gametocytes. Scale bar = 2 µm. **(B)** The progression of gametogenesis was examined in PfCAP-H^DiCre^ parasites, treated with rapamycin or DMSO, from Day 8 to Day 12 post-induction, by calculating different stages (Stage I to Stage V) with Hemacolor stained thin smears. The aberrant category includes parasites that appear stressed, distorted, and abnormal parasites. In contrast, the unknown category comprises the gametocytes, which were difficult to categorize based on standard categorization by *Carter et al., 1979*. The experiment was performed in two independent biological replicates. The error bar indicates SEM calculated by GraphPad Prism.

## Discussion

To unravel the function of PfCAP-H during the erythrocytic development in *Plasmodium falciparum*, we employed an inducible knockout parasite strain and ultrastructure expansion microscopy. We revealed that PfCAP-H is essential for asexual blood stages. The PfCAP-H knockout exhibited robust nuclear division defect, while the cytokinesis continued relatively normally. However, this is distinct from most eukaryotes, where the surveillance system verifies that nuclear division is completed typically before the onset of cytokinesis [38, 39]. Thus, this work corroborates previous studies that showed that karyokinesis and cytokinesis are independent, and these malarial parasites lack some aspects of the surveillance systems present in other eukaryotes [29, 40, 41].

In many eukaryotes, condensation of chromosomes during mitosis is a crucial step to ensure the faithful division of the genomic material in daughter cells. However, it is intriguing to comprehend how this equal distribution of genome is maintained and regulated in lower eukaryotes that possess decondensed forms of chromosomes throughout their life cycle [42]. Remarkably, despite the absence of condensed chromosomes, these lower eukaryotes have retained the highly conserved condensin complexes, which facilitate condensation of chromosomes [11]. This forces us to ponder the significance of these complexes in such an unusual scenario. So far, the function of condensin complexes in *Saccharomyces cerevisiae* has been extensively studied to decipher the equal distribution of the genome with such atypical decondensed chromosomes during closed mitosis [27, 43–45]. These complexes are recently identified in *Plasmodium spp.*, which share decondensed chromosomes similar to fission yeast [2, 14, 16]. However, attributing to their small nuclear size (∼1 µm diameter) and underexplored molecular mediators during mitosis compared to model organisms [29], it is challenging to investigate the chromosome dynamics in *Plasmodium spp.* during mitosis. The foremost step in the quest of interrogating chromosome dynamics is to get detailed evidence of different mitotic stages in these parasites. Strikingly, the recent establishment of ultrastructure expansion microscopy techniques in these parasites has unlocked the feasibility of studying the nuclear-related processes in these parasites [46]. Leveraging U-ExM capability to expand the size of parasites, we captured highly resolved images of mitotic events in these parasites. Furthermore, we showed that the PfCAP-H is highly expressed at the metaphase plate and propose that PfCAP-H could be used as a marker for the metaphase plate to mark the mitotically active nucleus during schizogony. Consequently, subcellular localization of PfCAP-H at the metaphase plate, along with the cues from localization of the mitotic spindle, as in the yeast mitosis model [43], would further aid in advancing our knowledge on mitosis in malarial parasites.

This study demonstrates that PfCAP-H interacts with PfCAP-G and is a crucial member of the condensin I complex. The U-ExM revealed that PfCAP-G does not localize to the mitotic chromosome in the absence of PfCAP-H, suggesting that PfCAP-H is required to load PfCAP-G on the mitotic chromosome, similar to its homologs in *Drosophila* and humans [18, 26]. Furthermore, the sequence analysis of PfCAP-H showed that PfCAP-H contains highly conserved N and C terminal region, which are the sites of interaction for SMC2/4 in its homologs [11, 21], inferring that PfCAP-H might also bind to SMC2/4 via these highly conserved regions. In *P. berghei*, it was shown that PbCAP-H co-immunoprecipitates with SMC2/4 in late schizonts but not in early schizonts [14]. This information hints that SMC2/4 can bind to chromosomes alone; however, PfCAP-H is required for assembling the subunits to function as a condensin I holo-complex. We propose that PfCAP-H plays a vital role in the assembly of the condensin I complex on the mitotic chromosomes in *Plasmodium* parasites, comparable to its homolog in other systems[17]. Furthermore, the essentiality of this complex suggests that chromosome condensation does occur in these parasites during mitosis, albeit to a lesser extent than in higher eukaryotes.

PfCAP-H is dynamically localized during schizogony with peak expression during its growth phase, followed by diffuse localization at the end of the schizogony. This expression and localization resemble the similar pattern displayed by the condensin I complex in several other eukaryotes during mitosis [9]. In model organisms, condensin complex localization and activity are tightly regulated through phosphorylation by mitotic kinases [47]. For example, the mitotic kinase cdc2-cyclin B phosphorylates the non-SMC subunits of condensin in *Xenopus* and human cells to condense the chromosome [48]. Barren, the *Drosophila* homolog of PfCAP-H, recruitment to mitotic chromosomes is facilitated by phosphorylation by Aurora B kinase [49]. Notably, phosphoproteomic analysis in *Plasmodium falciparum* showed that all the subunits of the condensin I complex, including PfCAP-H, are phosphorylated during asexual blood stages [6, 50]. We thus speculate that the condensin I complex is likely regulated by phosphorylation in these parasites. However, further study is required to investigate the mechanisms for controlling condensin I activity in *Plasmodium falciparum*.

Interestingly, while scrutinizing the localization of PfCAP-H in blood stages, we observed that some schizonts exhibited unconventional schizogony where already duplicated nuclei restart the next round of mitosis before completing karyokinesis. This observation agrees with previous studies of asexual blood stage development of *P. falciparum* [7, 51]. This variation in schizogony has been observed in the mosquito midgut during sporozoite formation of *Plasmodium spp.* [52] and is described as schizogony with limited karyokinesis [4]. Conversely, the evidence of such variation in the asexual blood stage suggests that schizogony is somewhat fluid and is not restricted only to classic schizogony. However, the factors responsible for such flexibility in blood stages remain to be explored.

In summary, we have shown that PfCAP-H is essential for the asexual blood stage development. PfCAP-H is critical for assembling the condensin I complex on the mitotic chromosomes. The knockout of PfCAP-H results in agglomeration of nuclei, possibly due to improper chromosome segregation, and defective karyokinesis – while cytokinesis remains largely normal (Fig. 9). Furthermore, PfCAP-H is not required for centrosome duplication or mitotic spindle assembly during schizogony. Overall, this study sheds light on the function of the condensin I complex during mitosis in these parasites and emphases their importance in lower eukaryotes with primitive mitotic features of decondensed chromosomes.

**Figure 9:**
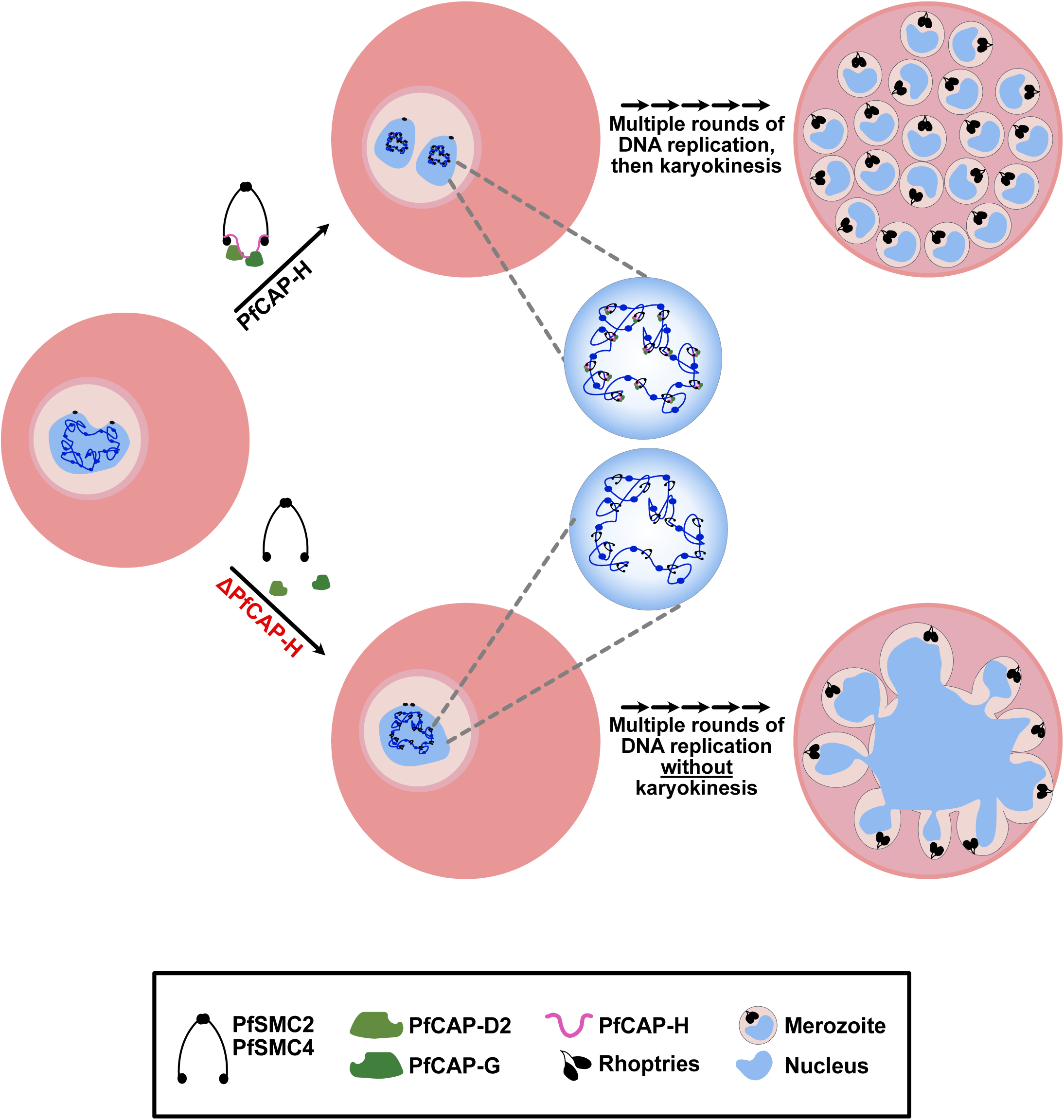
Proposed model of PfCAP-H function during blood stage development. Schematic representation of the proposed mechanism by which PfCAP-H function during schizogony. PfCAP-H plays a pivotal role in facilitating the formation of condensin I complex on the mitotic chromosomes. In parasites lacking PfCAP-H, the process of karyokinesis is significantly impaired, while the cytokinesis remains unaffected.

## Methods

### Plasmid construction

#### pPG25 (for smV5 tagged PfCAP-H)

The 5’ homology region (HR) and 3’ HR region were amplified from Pf3D7 genomic DNA with oJDD6763/6764 and oJDD6766/6767, respectively. The codon-altered PfCAP-H gene blocks (GB70.2 and GB71, IDT DNA) were amplified with oJDD6598/6599 and oJDD6196/6197, respectively. The smV5 and hDHFR region was amplified from pPG12 with oJDD6765/6086. The pGEM backbone for this plasmid was amplified from pPG12 with oJDD6083/6084. The six pieces were assembled with the Golden Gate *BsaI*-HF v2 assembly kit (NEB). Three guide RNA (gRNA) plasmids were used to target the PfCAP-H locus. The guides were annealed and ligated into *BpiI* digested pRR216 plasmid (SpCas9 expression plasmid) to construct pJPM59 (oJDD4921/4922), pCJM57 (oJDD5791/oJDD5792), and pCJM58 (oJDD5793/oJDD5794), respectively. The *EcoRI/NotI* linearized pPG25 plasmid was co-transfected with the three guide RNA-containing plasmids into 3D7*pfs47*DiCre parasites to generate PfCAP-H^DiCre^ parasites.

#### pRR85 (for generating BirA fused PfCAP-G)

The 5’ HR and 3’ HR regions were amplified from 3D7 genomic DNA with primers oJDD3881/3882 and oJDD3875/3876. Codon-altered PfCAP-G was amplified from gene block GB04 with oJDD3883/3884. The U6 promoter, gRNA for PfCAP-G, and U6 terminator were amplified from pBAM203 were assembled by overlapping PCR extension with oJDD3877/3878 and oJDD3879/3880. These five pieces were assembled by overlapping PCR and ligated into pRR28 cut with *NotI* and *XhoI*. The *SpeI* linearized pRR85 plasmid was co-transfected with a SpCas9 expression plasmid into the 3D7 parasite strain to generate PfCAP-G^BirA^ parasites.

#### pCJM22 (for generating smHA-tagged PfCAP-G)

The PfCAP-G region was digested from pRR85 with *NotI/XhoI*. The smHA and BSD sequences were obtained from pCJM01 by *NotI/XhoI* digestion. These two digested fragments were ligated with T4 DNA ligase to get pCJM22. For transfection, the Spe1 linearized pCJM22 plasmid was transfected with the SpCas9 expression plasmid in PfCAP-H^DiCre^ strain to generate PfCAP-H^DiCre^/PfCAP-G parasites.

#### pPG89 (for generating smHA-tagged PfNDC80)

The PfNDC80 region was PCR amplified with primers oJDD8427/8430 and digested with *NotI/XhoI*. The smHA and BSD sequences were obtained from pCJM22 by *NotI/XhoI* digestion. These two digested fragments were ligated with T4 DNA ligase to get pPG89. For transfection, the Spe1 linearized pPG89 plasmid was transfected with three guide RNA-containing plasmids in PfCAP-H^DiCre^ strain to generate PfCAP-H^DiCre^/PfNDC80 parasites.

All primers and geneblocks are shown in Supplemental Table S2.

### Reagents and Antibodies

All the primers used in this study were purchased from Thermo Fisher, gene blocks were purchased from IDT DNA, and restriction enzymes were purchased from New England Biolabs. The primary antibodies used in this study are the following: mouse α-V5 (clone SV5-Pk2, Bio-Rad); rabbit α-V5 (ICL, RV5-45A-Z), rat α-hemagglutinin (HA, clone 3F10, Sigma); mouse α-tubulin (clone B-5-1-2, Sigma); rabbit α-Histone H3 (ab1971, Abcam); mouse α-Centrin (CrCen clone 20H5, EMD Millipore); and rabbit α-PfGAP45 (gift from Julian Rayner at the University of Cambridge [53]). Secondary antibodies and other reagents (Alexa Fluor 405 NHS-Ester, SYTOX Deep Red Nucleic acid stain, and Hoechst 33342 solution) used for microscopy were purchased from Thermo Fisher.

### Sequence Alignments

The FASTA sequence of *Plasmodium falciparum* PfCAP-H (PF3D7_1304000) and its orthologs including *Plasmodium berghei* PbCAP-H (PBANKA_1402500), *Saccharomyces cerevisiae* BRN1 (P38170), *Schizosaccharomyces pombe* CND2 (Q9Y7R3) *Arabidopsis thaliana* CAPH (Q564K3), *Xenopus laevis* NCAP-H (O13067), *Homo sapiens* NCAP-H (Q15003), and *Drosophila melanogaster* Barren (P91663) were obtained from UniProt or PlasmoDB. The sequence identity and similarity was calculated using EMBOSS Needle [54]. Multiple sequence alignment was carried out using the default MUSCLE algorithm with MEGA software and aligned sequences were analyzed using ESPript 3.0 [55].

### *Plasmodium falciparum* culture and transfection

The *Plasmodium falciparum* 3D7 strain was obtained from the Walter and Elizabeth Hall Institute, and the 3D7*pfs47*DiCre parasite strain was obtained from Dr. Ellen Knuepfer [54]. Parasites were cultured in human O+ erythrocytes (deidentified from commercial vendor) at 4% hematocrit (HCT) in RPMI-1640 (Sigma) supplemented with 25 mM HEPES (4-(2-hydroxyethel)-1-piperazineethanesulfonic acid) (EMD Biosciences), 0.21% sodium bicarbonate (Sigma), 50 mg/l hypoxanthine (Sigma), and 0.5% Albumax II (Invitrogen) and were kept at 37 °C with a mixture of gases (5% CO_2_, 1% O_2_, and 95% N_2)_. Parasite growth was monitored by staining with Hemacolor (Sigma) staining solutions and observed under the microscope. The cultures were synchronized by combining Percoll and sorbitol treatments [56, 57].

To generate transgenic parasites, 60 μg of donor plasmid was linearized by digestion and co-transfected with 60 μg of SpCas9-plasmids containing gRNA into 10% ring-staged *P. falciparum* (3D7*pfs47*DiCre or 3D7) parasites by electroporation. The electroporation was performed at settings of 310 V, 950 μF and infinite Ω in a 0.2 cm cuvette with BioRad GenePulser. The parasites were cultured and selected with appropriate drugs. The PfCAP-H^DiCre^ and PfCAP-G^BirA^ parasites were selected with 2.5 nM WR99210 (Jacobus Pharmaceuticals), and PfCAP-H^DiCre^/CAP-G parasites was selected with 2.5 ug/ml blasticidin (Research Products International). Single clones of these transfected parasites were obtained by limitation dilution. The transgenic parasites were verified by PCR amplification using primers and/or whole genome sequencing of harvested genomic DNA (Biobasic blood genomic DNA miniprep kit). To induce excision of PfCAP-H gene, 0 – 4 h young parasites were treated with 100 nM rapamycin for up to 12 hrs, followed by washing with new culture media [28].

### Gametocyte induction

Trophozoite-stage parasites at 3% parasitemia and 2 % HCT were cultured with 50 % conditioned AlbuMax II medium. After 2 days, the culture was treated with 0.25 mg/ml heparin (Alpha Aesar A16198) and 2.5 mM N-acetylglucosamine to prevent subsequent invasion of asexual stage parasites. Gametocyte conversion and morphology was evaluated by blood smear stained with Hemacolor staining solution under light microscopy. The experiments were performed in two independent biological replicates. The gametocyte conversion was calculated by the ratio of gametocytes on day 6 to ring parasitemia on day 2. IFAs was performed on day 8 to localize the protein in gametocyte stages. The gametocytes were categorized as described [58].

### Replication assay

Synchronized ring-stage parasites were seeded with an initial parasitemia at 0.25 % in 1 % HCT, treated with DMSO or 20 nM rapamycin, and cultured for two consecutive cycles.100 μL culture was collected from each well on three different days (0, 2, and 4) and then washed once with 0.5% bovine serum albumin (BSA) in 1x PBS. The samples were resuspended in 100 μL staining solution containing 1:1000 SYBR Green I (Invitrogen) dilution in 0.5% BSA/PBS and incubated for 20 mins at RT. The stained samples were washed with 0.5 % BSA/PBS and then resuspended in 1x PBS. The number of infected red blood cells was determined by flow cytometry (FACSCalibur) using the CellQuest Pro program. The data from 100,000 cells was analyzed by FlowJo X and GraphPad Prism 9 software and represented as mean and SD of triplicates.

### Immunofluorescence assays (IFA)

For IFA, parasites were smeared onto slides and air-dried. The entire process of IFA was done in a humid chamber. The parasites were fixed with 4% (v/v) paraformaldehyde (PFA) in 1x PBS for 10 mins and rinsed quickly with 1x PBS. The fixed parasites were permeabilized with 0.1% Triton X-100 in PBS for 10 mins at room temperature (RT) and then washed three times with 1x PBS for 3 mins. The parasites were blocked with a 3% (w/v) BSA in PBS for 1h at RT. The smear was stained with respective primary antibodies (mouse α-V5 1:500, rat α-HA 1:250, mouse α-Centrin 1:500, rabbit α-V5 1:500) for 1h at RT or overnight at 4°C, followed by washing three times with 1x PBS for 5 mins. The samples were incubated with fluorescently labeled secondary antibodies (1:1000) for 30-45 mins at RT and washed thrice with 1x PBS for 5 mins to remove excess unbound antibodies. The DNA content of the parasites was stained with Hoechst 33342 (1:5000) in 1x PBS for 20 mins at RT and then quickly rinsed with 1x PBS. The parasites were mounted in Vectashield Vibrance antifade mounting media (Vector Laboratories Inc. H-1700) with coverslips and stored at 4°C until imaging. The z-stacked Images were acquired on a Zeiss LSM900 microscope with Airyscan 2 with 63X objective and analyzed using FIJI software.

### Ultrastructure Expansion microscopy (U-ExM)

For expansion microscopy, we followed a four-day protocol. On day 1, synchronized parasites at 4-5% parasitemia were collected and allowed to settle on a poly-D-lysine coated coverslip for 20 mins at 37 °C. Parasites were fixed with pre-warmed 4% (v/v) PFA for 20 mins at 37 °C, washed thrice with pre-warmed 1x PBS, and crosslinked with 1.4% formaldehyde and 2% acrylamide solution in 1x PBS overnight at 37 °C. On day 2, the gel polymerization was done in a gelation chamber. The parasite-coated coverslips were placed onto the mixture of TEMED/APS/monomer solution, kept on ice for 5 mins, and then incubated for 1 h at 37 °C. Post incubation, the coverslips plus gel were placed in a 6-well dish with 1ml denaturation buffer for 15 mins with agitation to displace gel from the coverslip. The detached gel was then incubated with 1.5ml denaturation buffer in a microcentrifuge tube for 90 mins at 95 °C. After cooling off, gels were transferred to a petri-dish containing 25ml ddH_2_O and incubated for 30 mins at RT. The gels were further incubated with 25 ml ddH_2_O overnight at RT to perform the first round of expansion. On day 3, gels were washed twice with 1x PBS for 15 mins at RT. The gels were blocked with 3% BSA/PBS for 30 mins at RT, followed by staining with respective primary antibodies (mouse α-V5 1:250, rat α-HA 1:100, mouse α-tubulin 1:500, mouse α-Centrin 1:250, rabbit α-GAP45 1:2500) in 3% BSA/PBS overnight at 4°C. On day 4, gels were washed thrice with 1x PBS with 0.5% Tween-20 for 10 mins at RT with agitation. Gels were stained with their respective secondary antibodies (1:500), Alexa Fluor 405 NHS-Ester (1:250), and SYTOX Deep Red dye nucleic acid stain (1:1000) in 1x PBS for 2h 30mins at RT protected from light. The stained gels were washed thrice with 1x PBS/0.5% Tween-20 for 10 mins at RT and then incubated with ddH_2_O for 30 mins. Water was replaced, and gels were incubated overnight at RT for the second round of expansion. The images were captured on a Zeiss LSM900 microscope with Airyscan 2 and analyzed using FIJI software.

### Immunoblots

Parasites were harvested by 0.02% saponin in PBS with protease inhibitors (PIC) (SigmaFast Protease Inhibitor Cocktail) and boiled in 1X Laemmli buffer supplemented with 1x PIC for 5 mins at 95 °C. The protein lysate (equivalent to 10^8^ parasites per lane) were run on a 4-20% Tris-glycine-sodium dodecyl sulfate gel and transferred to a PVDF membrane. The membrane was blocked with Licor Odyssey blocking buffer for 1 hr at RT. The immunoblot was probed with primary antibodies (α-V5 1:1000, α-HA 1:1000, and α-H3 1:2500), followed by incubation with secondary antibodies (1:1000) diluted in the Licor Odyssey blocking buffer. The immunoblot was scanned on a Licor Odyssey CLx imager system and quantified using volumetric measurement of fluorescence intensity with LiCor Image Studio 4.0.

### BirA Biotin Proximity-labeling

Synchronized 44-46 hour schizont-stage parasites (PfCAP-G^BirA^ and 3D7) were incubated with 150 μM biotin for 6 hours; the Protein Kinase G inhibitor, compound 1, was added to prevent parasite egress[59]. After this incubation, parasites were harvested with 0.05% saponin in 1x PBS and protease inhibitors. Parasite pellets were resuspended in 1ml of RIPA lysis buffer with PIC (50 mM Tris-HCl, Ph 7.5, 150 mM NaCl, 1% NP-40 (Tergitol), 0.5% Sodium deoxycholate, 0.1% Sodium dodecyl sulfate) for 1 h at RT on the rotator and sonicated with microtip sonicator with 30 s at 20 amplitudes, followed by 3 mins incubation on ice. This cycle was repeated twice. The lysate was spun down for 30 mins at max speed to remove hemozoin and other insoluble debris. The streptavidin-coated beads were washed with RIPA/PIC and incubated with the cleared lysate overnight on the rotator at 4°C. After incubation, beads were sequentially washed with RIPA buffer and Wash Buffer 1 (2% SDS), Wash buffer 2 (50 mM HEPES, pH 7.5, 500 mM NaCl, 1 mM EDTA, 1% Triton-X 100, 0.1% Sodium deoxycholate), Wash buffer 3 (10 mM Tris HCl, pH 8.0, 250 mM Lithium Chloride, 1 mM EDTA, 0.5% NP-40, 0.5% Sodium deoxycholate), and Wash buffer 4 (50mM Ammonium bicarbonate in ddH20). The sample was resuspended in 40 μL of wash buffer 4 and stored at -80 °C until further processing for mass spectrometry. On-bead digestion, followed by LC/MS-MS and data analysis, was performed at the Harvard Taplin Mass Spectrometry Facility. The results were analyzed by comparing the unique and total peptides between PfCAP-G^BirA^ and 3D7 parasites in three independent biological replicates.

## Acknowledgements

We acknowledge the support of Ross Tomaino at the Taplin Mass Spectrometry Facility and Paula Montero-Llopis of the Harvard Medical School MicRoN core. This work was supported by the National Institutes of Health R01 AI145941 (J.D.D).

## Author Contributions

P.G.: Conceptualization, Methodology, Formal Analysis, Investigation, Writing–Original Draft Preparation, Visualization. J.P.M.: BioID experiment and analysis. J.D.D.: Conceptualization, Data Curation, Supervision, Project Administration, Funding Acquisition, Writing–Review and Editing.

The authors declare no competing interests.

## Data Availability

All data generated throughout this study are incorporated into the manuscript and supplementary files. Whole genome sequencing data have been deposited in the NCBI Sequence Read Archive (#XXXXX). Protocols, raw data, or any materials employed in this study are available upon request.

