## Supplemental Figures for "PfCAP-H is essential for assembly of condensin I complex and karyokinesis during asexual proliferation of *Plasmodium falciparum*"

**Supplemental Figures and Legends**

**A**

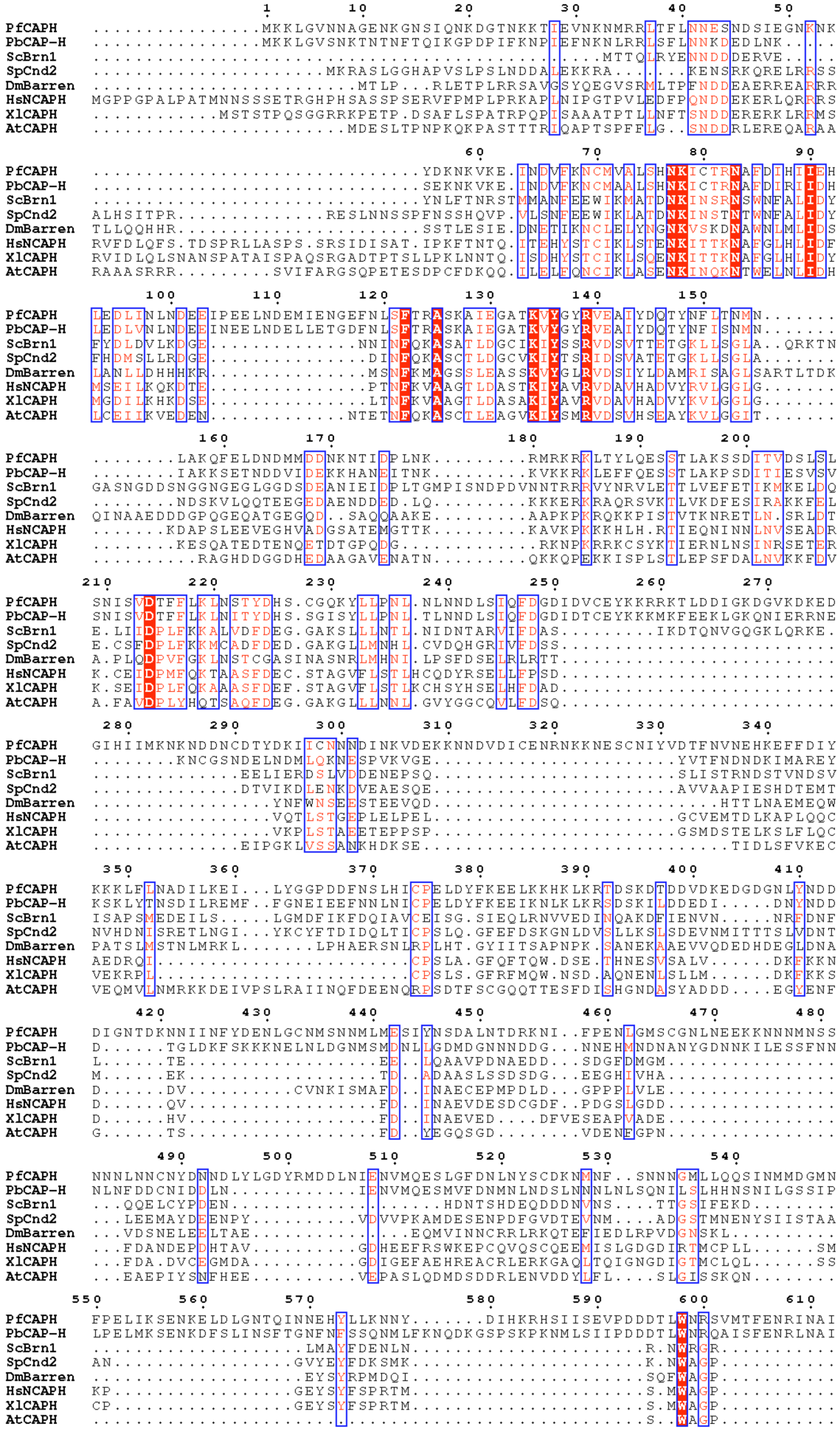
PfCAPH
PbCAP-H
ScBrn1
SpCnd2
DmBarren
HsNCAPH
XlCAPH
AtCAPH

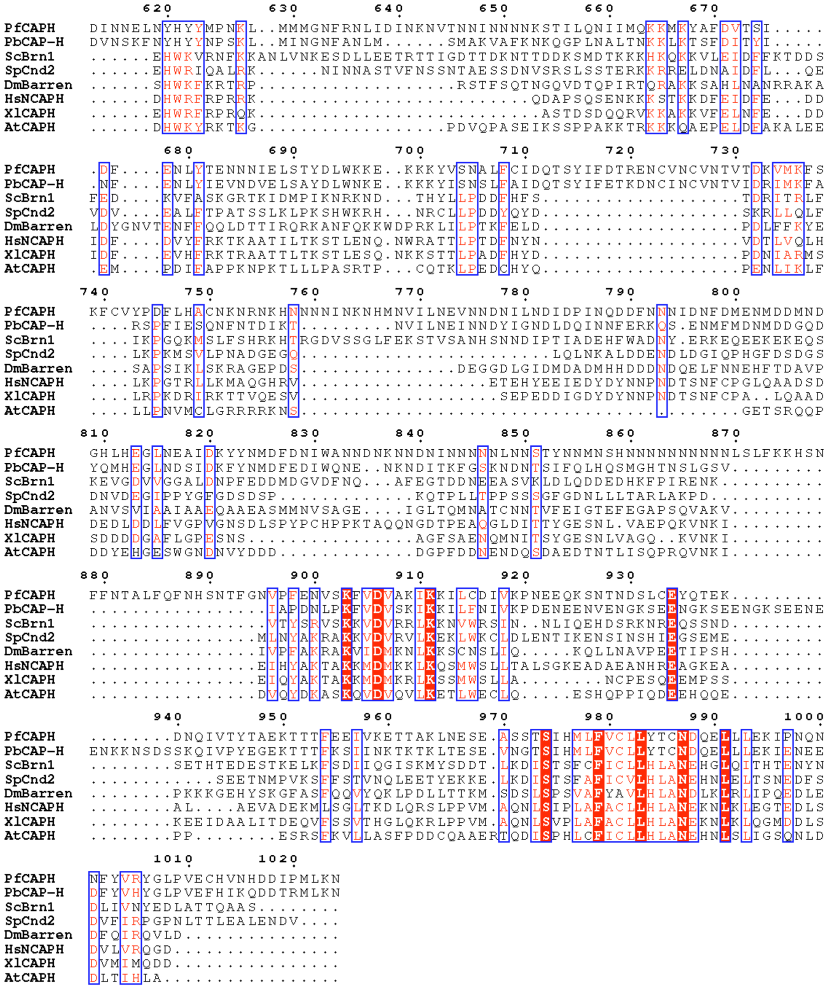

**B**

| | PbCAP-H | ScBrn1 | SpCnd2 | DmBarren | HsNCAPH | XlCAPH | AtCAPH |
|---|---|---|---|---|---|---|---|
| **Similarity** | 64.8 | 31.5 | 33.8 | 21.2 | 26.1 | 22.2 | 29.1 |
| **Identity** | 49.7 | 16.5 | 18 | 12 | 12.6 | 12.3 | 15.7 |

**Fig. S1: In silico analysis of PfCAP-H and its homologs**. **(A)** Sequences of *P. falciparum* PfCAP-H (PF3D7_1304000) and its homologs, including *P. berghei* PbCAP-H (PBANKA_1402500), *Saccharomyces cerevisiae* BRN1 (P38170), *Schizosaccharomyces pombe* CND2 (Q9Y7R3), *Arabidopsis thaliana* CAPH (Q564K3), *Xenopus laevis* NCAP-H (O13067), *Homo sapiens* NCAP-H (Q15003), and *Drosophila melanogaster* Barren (P91663) were aligned using default MUSCLE algorithm (MEGA 11 software) and analyzed using ESPript 3.0 [52]. **(B)** The table comprises the sequence identity and similarity calculated and compared for PfCAP-H and its homologs using EMBOSS Needle [51].

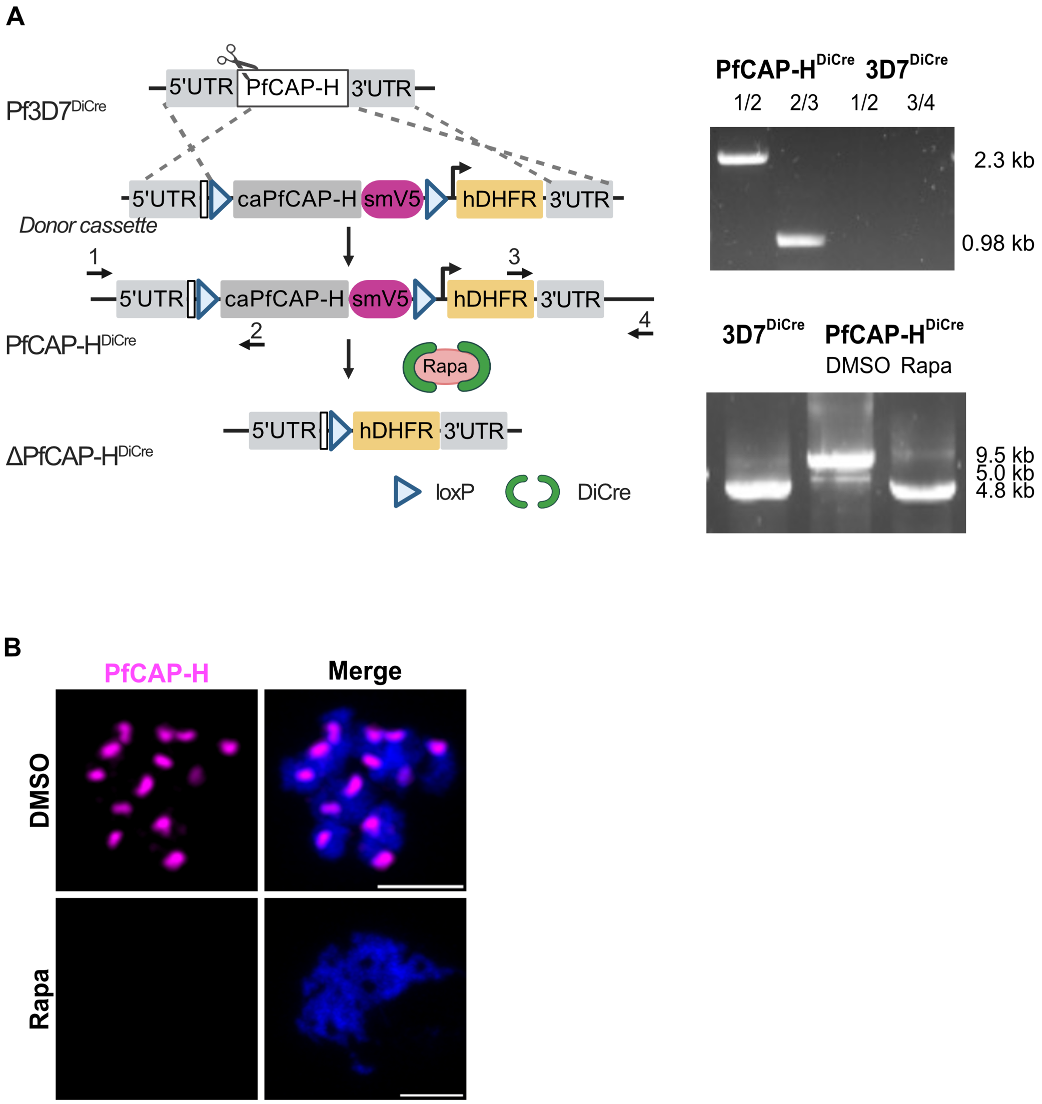


**Fig. S2: Generation of inducible knockout PfCAP-H$^{DiCre}$ parasite strain. (A)** Schematic of strategy to generate PfCAP-H$^{DiCre}$ parasite. In PfCAP-H$^{DiCre}$ parasites, we replaced the WT endogenous locus of PfCAP-H in Pf3D7$^{DiCre}$ parasites by a donor cassette containing codon-altered PfCAP-H with spaghetti monster (sm)-V5 tag (magenta-colored box) at the C-terminal flanked by two loxP sites (triangle) and hDHFR (orange-colored box) as a selectable marker in Pf3D7$^{DiCre}$ parasites. This schematic is also show in Figure 1 and reproduced here for clarity of the PCR integration checks. Top DNA gel: the modified locus was verified by integration PCR amplification with primers 1 and 2 (1/2) [2.3 kb, (primer 1 is oJDD5046 and primer 2 is oJDD6315)] and primers 3 and 4 (3/4) [0.98 kb, (primer 3 is oJDD3690 and primer 4 is oJDD5223)]. Bottom DNA gel: PCR amplification of entire locus with primers 1 and 4 in parental parasite, and PfCAP-H$^{DiCre}$ after treatment with rapamycin or DMSO vehicle control. The parental locus is 4.8 kb, the transgenic locus is 9.5 kb, and the excised transgenic locus is 5.0 kb. **(B)** Schizonts from rapamycin/DMSO treated PfCAP-H$^{DiCre}$ parasites were fixed and stained with $\alpha$-V5 against PfCAP-H and Hoechst *33342* for DNA. The slide-based IFA showed that PfCAP-H is absent following rapamycin treatment. Scale bar = 2 μm.

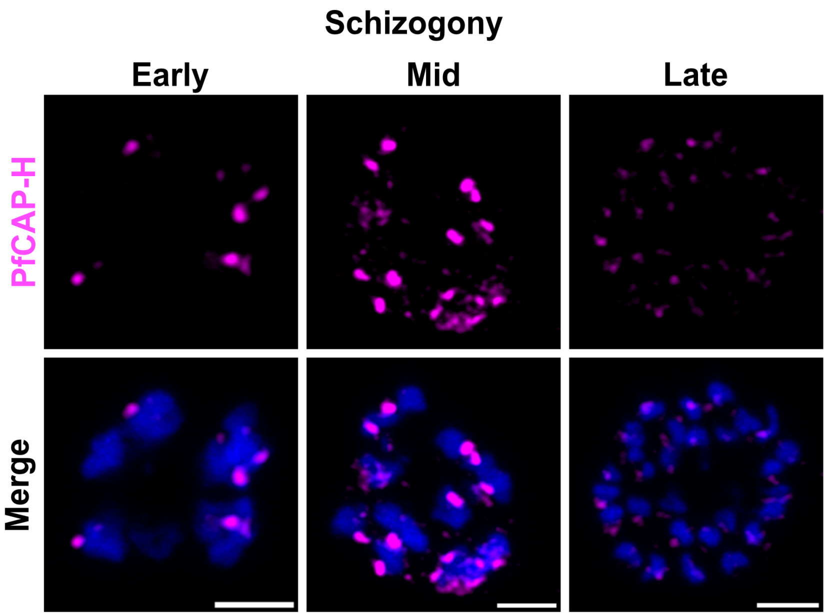


**Fig. S3: Expression of PfCAP-H during schizogony.** The expression of PfCAP-H during schizogony was visualized by α-V5 (magenta) for smV5 tagged PfCAP-H by slide-based IFA. The IFA demonstrates that PfCAP-H is strongly expressed during early- and mid-schizogony. In late schizonts, the PfCAP-H expression is diminished. The DNA was stained with Hoechst *33342* (blue). Scale bar = 2 μm.

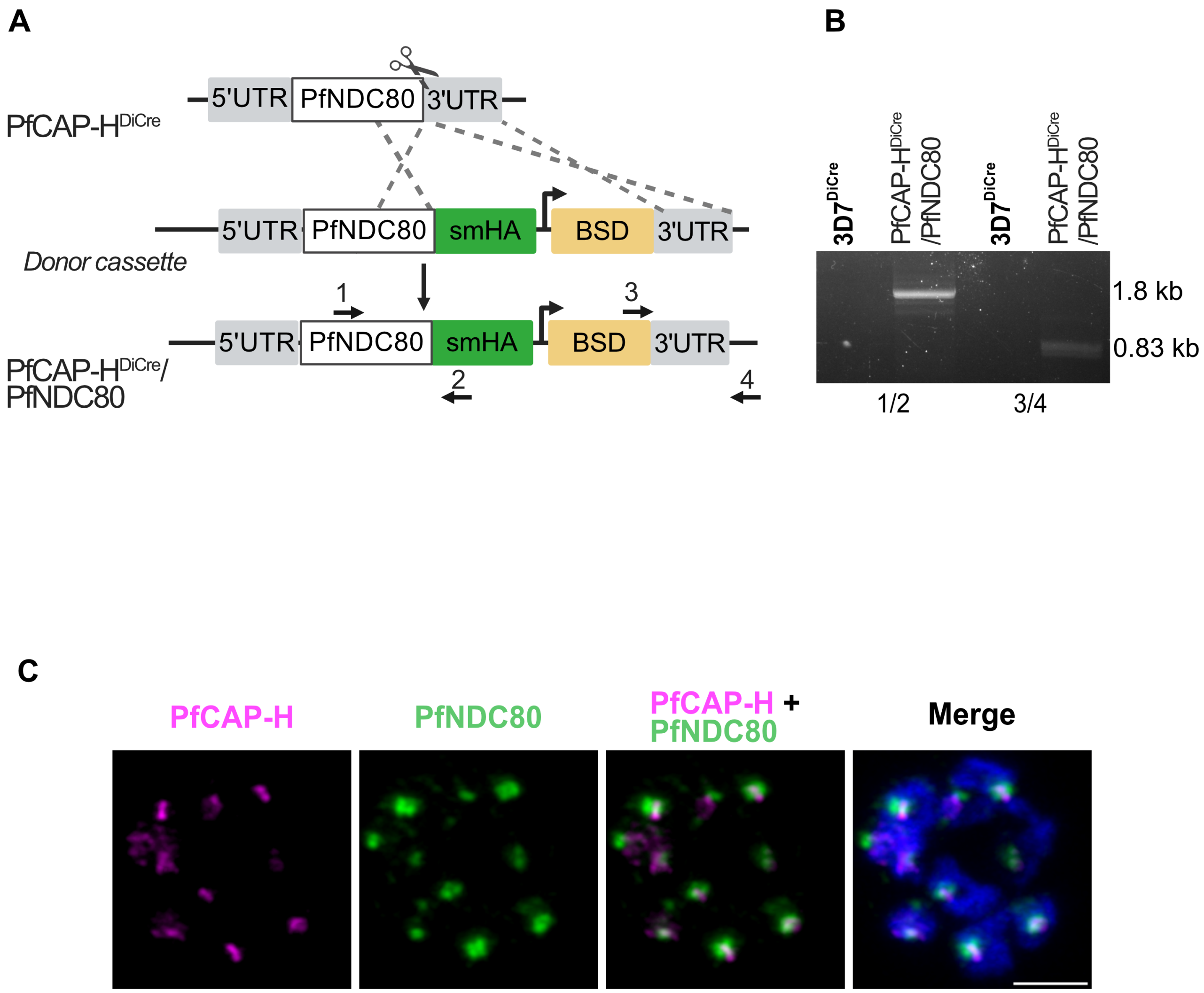


**Fig. S3: Generation of PfCAP-H$^{DiCre}$/PfNDC80 parasite strain. (A)** Schematic of strategy to generate PfCAP-H$^{DiCre}$/PfNDC80 parasite by Cas9 gene editing. **(B)** In PfCAP-H$^{DiCre}$/PfNDC80 parasites, we introduced a smHA tag (green-colored box) at the C-terminal end of the WT endogenous locus of PfNDC80 and Blasticidin-S-deaminase (BSD, yellow-colored box) as a selectable marker. The modified locus was verified for integration by PCR amplification with primers 1 and 2 (1/2) [1.8 kb, primer 1 is oJDD8542 and primer 2 is oJDD733)] and primers 3 and 4 (3/4) [0.83 kb, primer 3 is oJDD6312 and primer 4 is oJDD8543)]. **(C)** The expression of PfCAP-H ($\alpha$-V5, magenta) and PfNDC80 ($\alpha$-HA, green) was visualized with respective antibodies at schizont stage by slide-based IFA. The DNA was stained with Hoechst *33342* (blue). Scale bar = 2 μm.

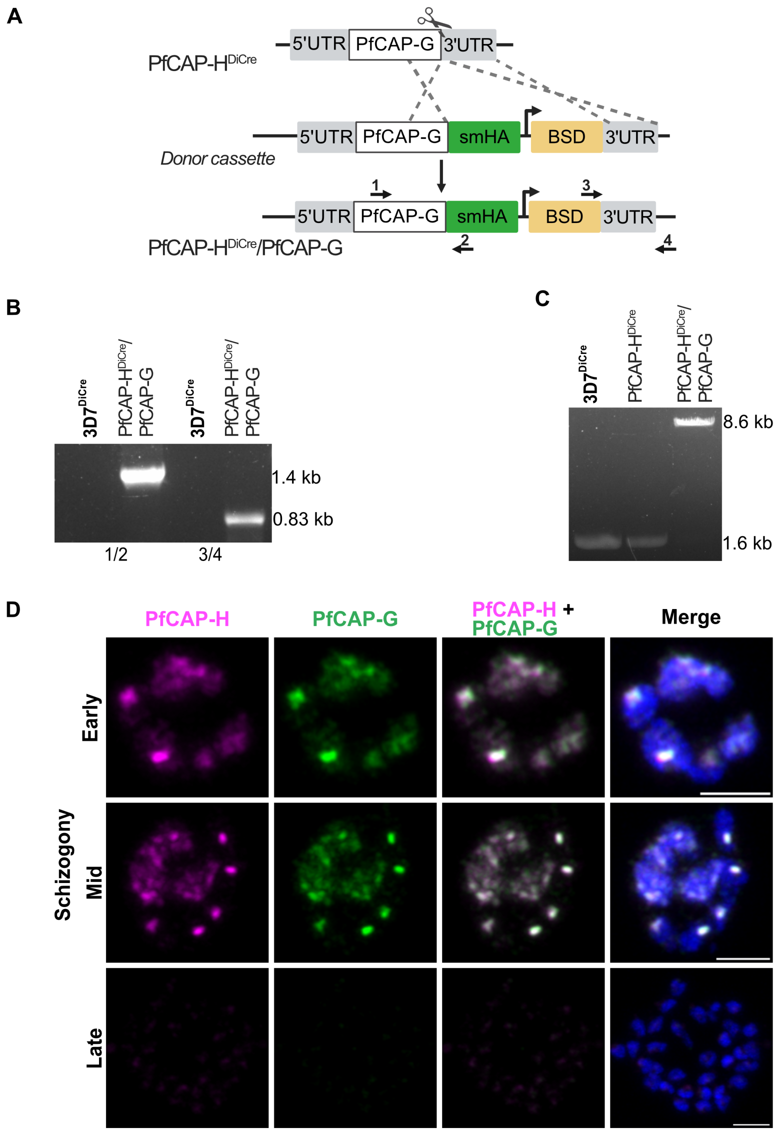


**Fig. S5: Generation of PfCAP-H$^{Dicre}$/PfCAP-G parasite strain and expression of PfCAP-H and PfCAP-G during schizogony**. **(A)** Schematic of strategy to generate PfCAP-H$^{DiCre}$/PfCAP-G parasite by Cas9 gene editing. In PfCAP-H$^{DiCre}$/PfCAP-G parasites, we introduced smHA tag (green-colored box) at the C-terminal of the WT endogenous locus of PfCAP-G and Blasticidin (BSD, yellow-colored box) as a selectable marker. **(B)** The modified locus was verified for integration by PCR amplification with primers 1 and 2 (1/2) [1.4 kb, primer 1 is oJDD7889 and primer 2 is oJDD2934] and primers 3 and 4 (3/4) [0.83 kb, primer 3 is oJDD4971 and primer 4 is 7890] regions. **(C)** Furthermore, the PCR amplification using primers 1 and 4 showed that the PfCAP-G was modified only in PfCAP-H$^{DiCre}$/PfCAP-G as compared to PfCAP-H$^{DiCre}$ and Pf3D7$^{DiCre}$ parental parasite strains. **(D)** The expression of PfCAP-H ($\alpha$-V5, magenta) and PfCAP-G ($\alpha$-HA, green) was visualized with respective antibodies during schizogony by slide-based IFA. The IFA showed that PfCAP-H is expressed brightly during early and mid-schizogony while the expression is diminished in late segmentation. PfCAP-G also showed similar expression pattern as PfCAP-H. The DNA was stained with Hoechst *33342* (blue). Scale bar = 2 µm.

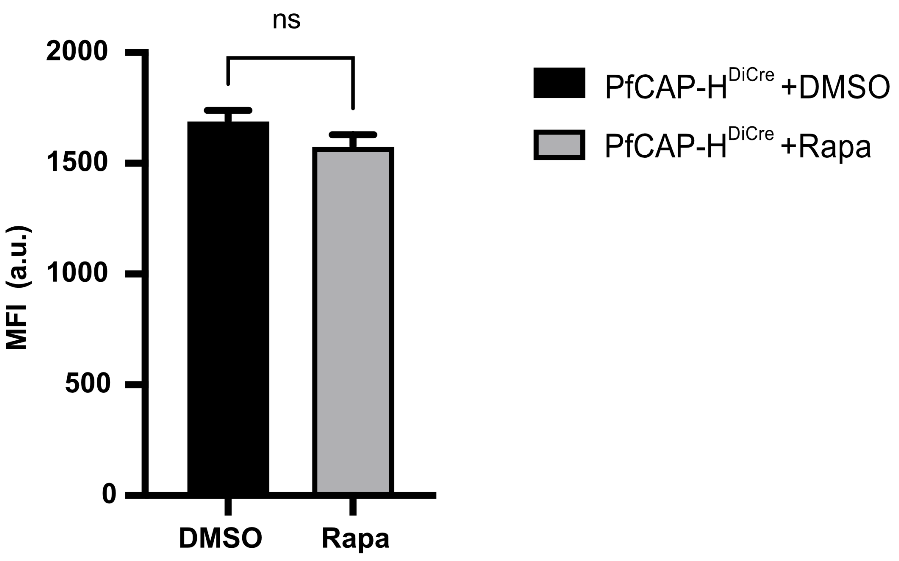


**Fig. S6: Effect of PfCAP-H knockout in DNA replication during blood stages of PfCAP-H$^{DiCre}$ parasites**. The PfCAP-H$^{DiCre}$ parasites were treated with rapamycin/DMSO at 40 hpi. The DNA content was measured by quantifying the mean fluorescence intensity (MFI) of SYBR Green stained parasites with flow cytometry. The analysis did not find any significant difference in DNA content between DMSO and rapamycin treated parasites. The experiment was done in triplicate and data were analyzed using GraphPad Prism.

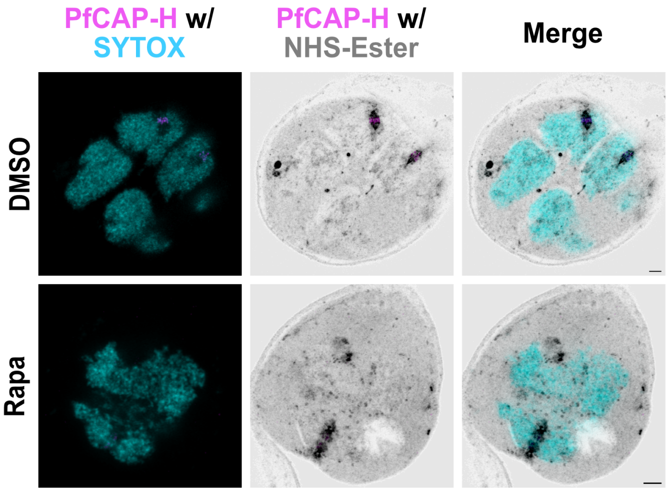


**Fig. S7: Effect of PfCAP-H knockout on karyokinesis.** Synchronized PfCAP-H$^{DiCre}$ parasites were treated with rapamycin or DMSO, probed with α-V5 (PfCAP-H, magenta), NHS-Ester (protein, greyscale), and SYTOX (nucleus, cyan) for U-ExM (n= 3). Loss of PfCAP-H results in incompletely separated nuclei in early schizonts. Scale bar = 2 μm.

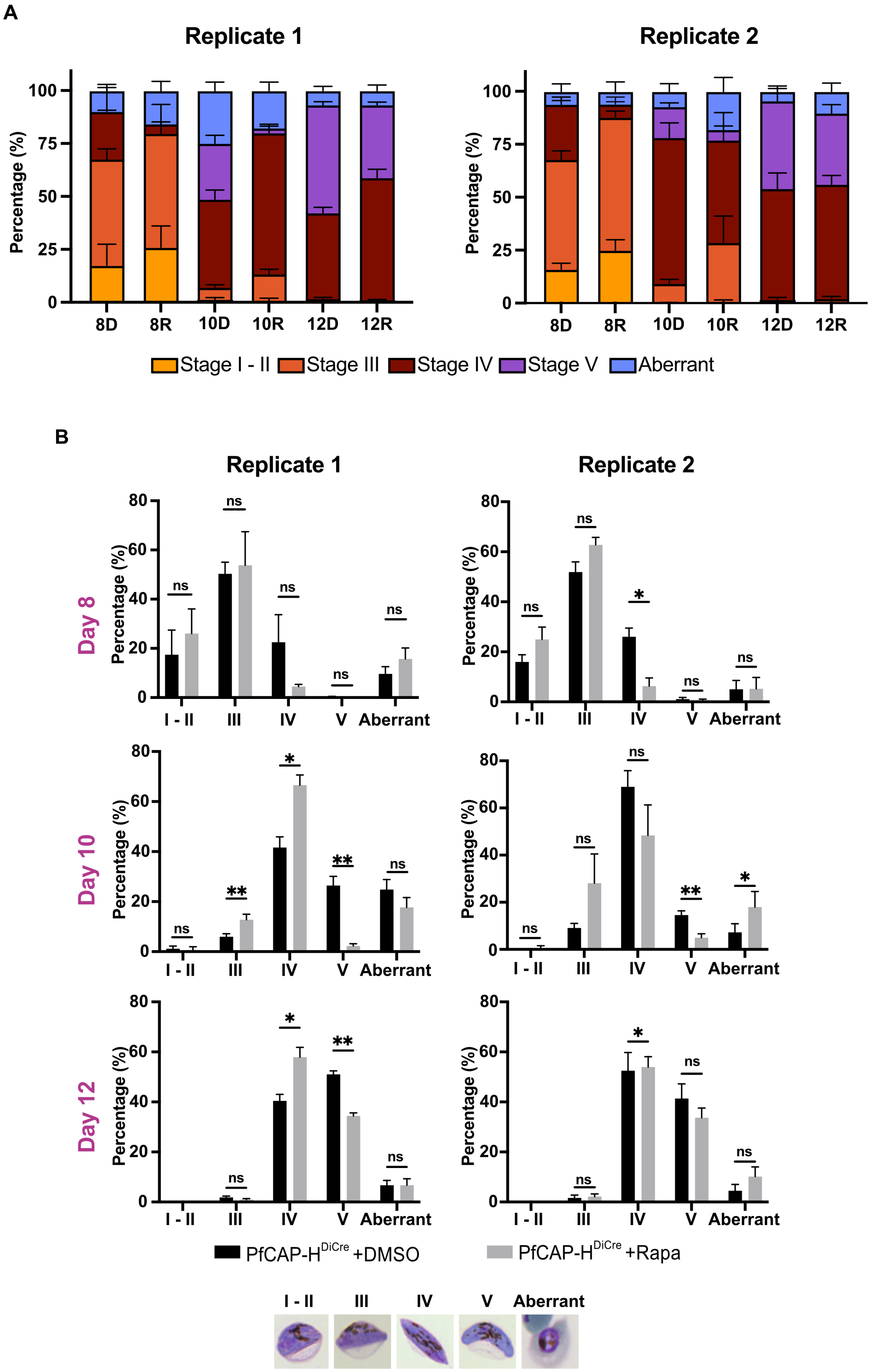


**Fig. S8: Effect of PfCAP-H KO on gametocytogenesis. (A and B)** The progression of gametogenesis was examined in PfCAP-H$^{DiCre}$ parasites, treated with rapamycin or DMSO, from Day 8 to Day 12 post-induction, by calculating different stages (Stage I to Stage V) with Hemacolor stained thin smears. The aberrant category includes parasites that appear stressed, distorted, and abnormal parasites. These stages were categorize based on standard criteria by *Carter et al., 1979*. The experiment was performed in two independent biological replicates. The error bar indicates SEM calculated by GraphPad Prism.
